# Hyperspherical geometry positions the lipidome as a partly independent axis of human brain organization

**DOI:** 10.1101/2025.09.18.677162

**Authors:** Maria Osetrova, Arsenii Onuchin, Elena Stekolshchikova, Philipp Khaitovich, Kirill Polovnikov

## Abstract

The brain’s transcriptome is well mapped, but the spatial organization of lipids—over half of brain dry mass—remains poorly defined. We profiled lipidomic (419 species) and transcriptomic (15,013 genes) signatures from 35 anatomically defined regions in four healthy adult donors, measured from the same tissue samples. Both modalities recapitulate major neuroanatomical divisions, yet the lipidome shows distinctive features: a pronounced white–gray asymmetry, a smooth neocortical rostrocaudal gradient, and limbic-specific lipid clusters absent in transcriptomic space. Using regression against gene-expression principal components and two null models (random and anatomy-aware), we define three data-driven classes of lipid–gene relationships: *Synchronizers* (10%) tightly coupled to gene programs, *Anchors* (67%) predictable at levels expected from gross anatomy and cell-type composition, and *Drifters* (23%) largely transcription-independent. A simple geometric framework unifies these patterns: after centering and normalization, molecular profiles lie on a hypersphere where two interpretable coordinates—polar latitude relative to transcriptome-defined white/gray poles and nearest-gene angular distance—jointly index coupling; an analytical null quantitatively explains the observed scaling. Key effects are robust in leave-one-donor-out analyses and position the lipidome as a partly independent organizational axis of the human brain. Broadly, our results provide an interpretable geometric framework for multi-omics integration, supported by an interactive platform.

## Introduction

Mapping the human brain organization at molecular resolution is one of the central challenges of modern biology. Furthermore, the molecular organization may differ among the biochemical data types. Indeed, studies combining transcriptome, proteome, and metabolome analyses have revealed that these different molecular layers can exhibit strikingly different organizational principles within a single biological system [1, 2, 3, 4, 5, 6, 7, 8, 9, 10], from developmental gene-expression gradients to proteomic signatures of functional demand [11, 12, 13, 14, 15]. Integrating information across these layers has uncovered hidden regulatory modules and novel axes of brain organization that are invisible in single-modality studies [16, 17, 18, 19, 20, 21, 22, 23, 24]. Yet, despite rapid progress, a substantial section of molecular brain organization, the brain lipidome, remained largely absent from this effort.

Lipids are the brain’s most abundant molecular constituents, comprising over half of its dry mass [73, 74]. Lipids define membrane architecture, regulate synaptic signaling, and fuel metabolic processes [25]. Their distribution is sharply polarized: the brain’s white matter is lipid-rich, dominated by cholesterol and sphingolipids, whereas gray matter favors polyunsaturated species that enhance membrane fluidity [26]. Despite their ubiquity and multi-faceted functionality, lipids remain conspicuously absent from multi-omics brain atlases, with prior studies limited to fragmentary regional snapshots [26]. This gap of knowledge obscures a fundamental question: *Does the lipidome merely echo transcriptional programs, or does it constitute an autonomous organizational layer of the human brain?*

Here we address this question by integrating lipidomic and transcriptomic profiles across *n* = 35 anatomically defined regions of the healthy adult human brains. By combining network-based analyses with a hyperspherical geometric framework, we reveal shared and divergent organizational features of these two molecular phenotypes. While the broad picture of the transcriptome and lipidome variation across the brain aligns with its broad neuroanatomical divisions, the lipidome exhibits unique patterns, including a white-gray asymmetry, rostrocaudal gradients in the neocortex, and limbic-specific clustering absent in the transcriptome. Using regression modeling, we introduce three categories of lipid–gene relationships—synchronizers (10%, tightly gene-coupled), anchors (67%, anatomy-constrained), and drifters (23%, largely transcriptome-independent). Operationally, we benchmark lipid–gene predictability against both a random and an anatomy-aware null that preserves within-division composition; lipids exceeding only the random threshold are classified as anchors, reflecting anatomical/compositional constraints rather than gene-specific coupling. In this framework we do not treat regional composition as a nuisance regressor to be removed. Rather, composition is the anatomical baseline: Anchors are the molecules whose predictability matches this composition-preserving benchmark, while Synchronizers demonstrate coupling above it.

A simple geometry explains these classes: after centering and normalization, each molecular profile lies on a high-dimensional hypersphere, where two coordinates summarize coupling—latitude *θ* (position relative to the transcriptome-defined white pole) and the nearest-gene angular distance Δ (the geodesic angle to the closest transcript vector). We further derive an analytical null model of random gene placement predicting how nearest-gene distance scales with latitude 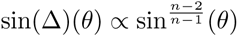, which quantitatively matches the data and explains why predictability declines away from the pole: *Synchronizers* cluster at low Δ(*θ*), *Anchors* at intermediate values, and *Drifters* near the equator with large Δ(*θ*). In summary, we establish the lipidome as a distinct, partly independent axis of human brain organization and present (*θ,* Δ) as an interpretable coordinate system that unifies multi-omics integration. We show that all central results are stable in leaveone-donor analyses.

To make these findings accessible, we provide an interactive platform (https://humanlipidome.pythonanywhere.com) for exploring inter-molecular maps on the hypersphere across brain regions.

## Results

### Integrated network organization of lipidome and transcriptome aligns with gross anatomy but reveals modality-specific differences

To compare lipidomic and transcriptomic organization of the human brain, we analyzed lipid (419 species) and mRNA (15,013 genes) profiles across 35 regions from four healthy adult donors [33, 71], with both modalities measured from the same tissue samples (Figure 1a; Tables S1–S3). These regions span five major anatomical divisions—cerebellar gray matter (1), white matter (4), neocortex (20), limbic system (5), and basal ganglia (5) (Table S2)—matching those used in previous transcriptomic atlases [23, 27, 28, 29] and ensuring broad coverage of distinct cytoarchitectural territories.

**Figure 1:**
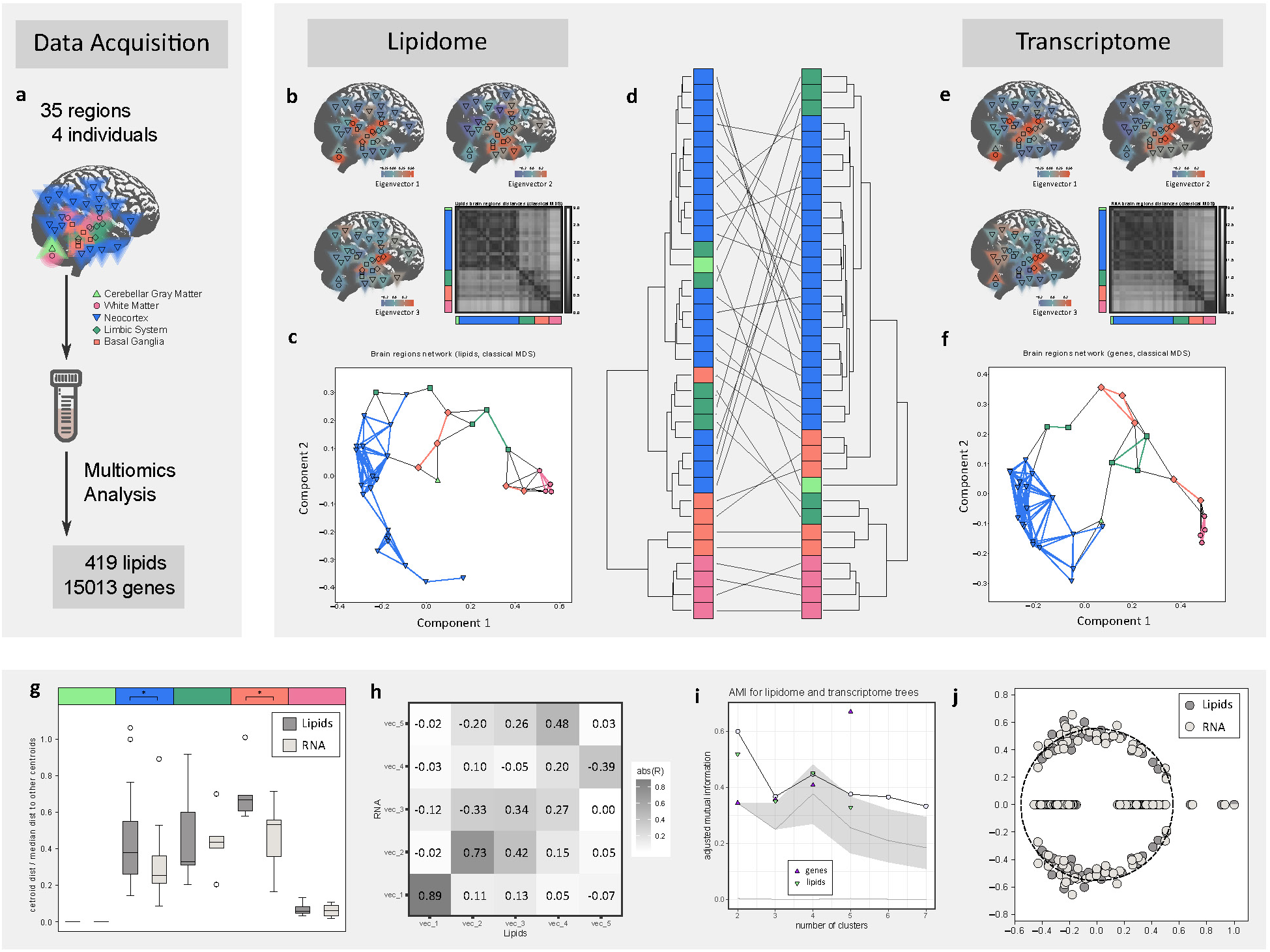
Comparative network organization of the brain lipidome and transcriptome. **a**, Anatomical map of the 35 sampled brain regions grouped into five major structures: cerebellar gray matter, white matter, neocortex, limbic system, and basal ganglia, with the number of unique lipids and genes identified per region. **b-c**, Classical multidimensional scaling (MDS) of the lipidome network based on pairwise regional distances, showing separation of brain systems. Colors reflect values of the first three eigenvectors of the Gram matrix. **d**, Hierarchical clustering of brain regions based on lipidomic (left) and transcriptomic (right) similarity. **e-f**, Equivalent MDS embedding of the transcriptome network. **g**, Evaluation of clustering quality for anatomical structures within lipidomic and transcriptomic embeddings. Lower values indicate better separation of gross anatomical divisions. **h**, Pairwise correlations of eigenvectors between lipidome- and transcriptome-derived embeddings. **i**, Adjusted mutual information (AMI) quantifying overlap between lipidomic and transcriptomic hierarchical clusterings compared to a null model (gray curve). Triangles mark AMI scores at increasing anatomical resolution (2–5 clusters). **j**, Eigenvalue spectrum of the Newman flow matrix, with isolated eigenvalues (right of dotted line) corresponding to community structure in both lipidome and transcriptome networks. Overall, lipidomic and transcriptomic maps both recapitulate major anatomical divisions, but their clustering structures only partially overlap, indicating complementary organizational principles.

We first constructed molecular similarity networks, treating each region as a node and connecting them by pairwise correlations of lipid or gene expression profiles (Methods). Dimensionality reduction consistently revealed strong correspondence between lipidomic and transcriptomic spaces. Classical multidimensional scaling (MDS) explained 33.4%, 14.7%, and 8.1% of variance in the lipidome and 30.1%, 10.1%, and 6.1% in the transcriptome across the first three components (Figure 1b,e). Cross-modal correlations of matching eigenvectors showed close alignment for the first two axes (*r*_11_ = 0.89*, r*_22_ = 0.73; Figure 1h), indicating shared large-scale topology.

In both modalities, the first component robustly captured the gray–white matter division, the dominant axis of organization. The second distinguished basal ganglia from cortical regions, while the third showed weaker specificity with partial separation of limbic areas (Figure 1b,e). Thus, both molecular layers are intrinsically structured by gross anatomy, with gray–white matter contrast emerging as a universal organizing principle.

Unsupervised clustering corroborated this pattern. Distance matrices derived from the first three MDS components revealed block-diagonal structures corresponding to anatomical subdivisions of white matter and neocortex (Figure 1b,d,e). Limbic and basal ganglia regions were more diffusely organized, with some affiliating to neocortex and others to white matter. Two-dimensional embeddings highlighted clear community boundaries, with neocortical and white matter regions consistently segregating into opposite hemispaces (Figure 1c,f). This stable dichotomy underscores the tight coupling between molecular architecture and gross anatomy. Despite these global similarities, modality-specific differences emerged. In the MDS embeddings, the neocortex cluster was clearly denser in transcriptomic space, with nearly twice as many intra-cluster edges (110 vs. 61) and 10% shorter average interregional distances relative to the lipidomic network (darker blocks in Figure 1b,e). To quantify these differences, we calculated cluster compactness as the ratio of intra-to inter-cluster Euclidean distances (Figure 1g). White matter regions formed the most distinct cluster in both modalities, with compactness differing by *<* 10%. In contrast, neocortical regions were significantly more compact in the transcriptomic network (Mann–Whitney U test, *p <* 0.05). Basal ganglia also clustered more tightly in transcriptomic space (*p <* 0.05), consistent with stronger transcriptional co-regulation, while limbic regions showed only a non-significant trend in the same direction (*p <* 0.1). Together, these results indicate that although both molecular networks reflect anatomical organization, transcriptomic patterns provide sharper resolution of neocortical and basal ganglia structure.

Hierarchical clustering analyses corroborated these findings while highlighting additional modality-specific distinctions. Using divisive hierarchical clustering—an unsupervised method that recursively partitions networks into nested clusters (Methods)—we observed minor but consistent differences in the placement of limbic, basal ganglia, and cerebellar regions between lipidomic and transcriptomic networks (Figure 1d). For example, the cerebellar cortex formed a distinct cluster in the transcriptomic but not in the lipidomic representation. Similarly, limbic regions diverged between modalities: in the transcriptomic network, the medial dorsal thalamus and hypothalamus clustered with white matter, whereas the entorhinal cortex, amygdala, and hippocampus aligned with neocortex in both modalities (Figure S1). Thus, while clustering and low-dimensional embeddings converge on a coherent large-scale framework, they also reveal representation-specific variation in the finer anatomical placement of subcortical and limbic structures.

Quantitative assessment using adjusted mutual information (AMI) [32] confirmed that both modalities contained significantly more structural information than random groupings (permutation test, *p <* 0.05 across 2–7 clusters, except at 4 clusters; Figure 1i). Importantly, lipidomic and transcriptomic clusterings agreed more closely with each other than with purely anatomical partitions, indicating complementary rather than redundant organization.

Finally, spectral analysis of non-backtracking flow matrices provided an objective estimate of intrinsic cluster number [30, 31]. Both networks yielded three significant eigenvalues beyond the bulk distribution (Figure 1j), consistent with a tripartite organization: (i) white matter, (ii) neocortex, and (iii) a transitional module comprising limbic and basal regions. This three-cluster architecture was shared across modalities despite the transcriptome’s finer resolution within neocortex and basal ganglia.

To assess robustness to inter-individual variation, we performed leave-one-donor-out (LODO) analyses: in four iterations, each donor was removed in turn and the full pipeline rerun. The dominant gray–white division persisted in both lipidomic and transcriptomic spaces, with highly similar MDS embeddings for the lipidome (Fig. S2) and transcriptome (Fig. S3). The principal axes also remained aligned across modalities: correlations between the top two lipidomic and transcriptomic eigenvectors were consistently high in all LODO runs (Fig. S4). Non-backtracking (flow) spectral analysis likewise recovered a stable tripartite community structure on every iteration—three isolated eigenvalues—corresponding to white matter, neocortex, and a transitional limbic/basal module (Fig. S5). Together, these tests indicate that the key organizational signals are not driven by any single donor.

### Laplacian eigenmaps reveal geometric encoding in molecular brain networks

The strong white–gray matter segregation observed in both modalities led us to ask whether finer-scale neuroanatomical geometry could also be recovered from molecular profiles. To address this, we applied Laplacian eigenmaps—a nonlinear dimensionality reduction method that preserves both local community structure and global geometry (Methods; Figure 2a). The leading eigenvectors of the normalized graph Laplacian provided interpretable coordinates for embedding brain regions in two dimensions for both transcriptomic and lipidomic networks.

**Figure 2:**
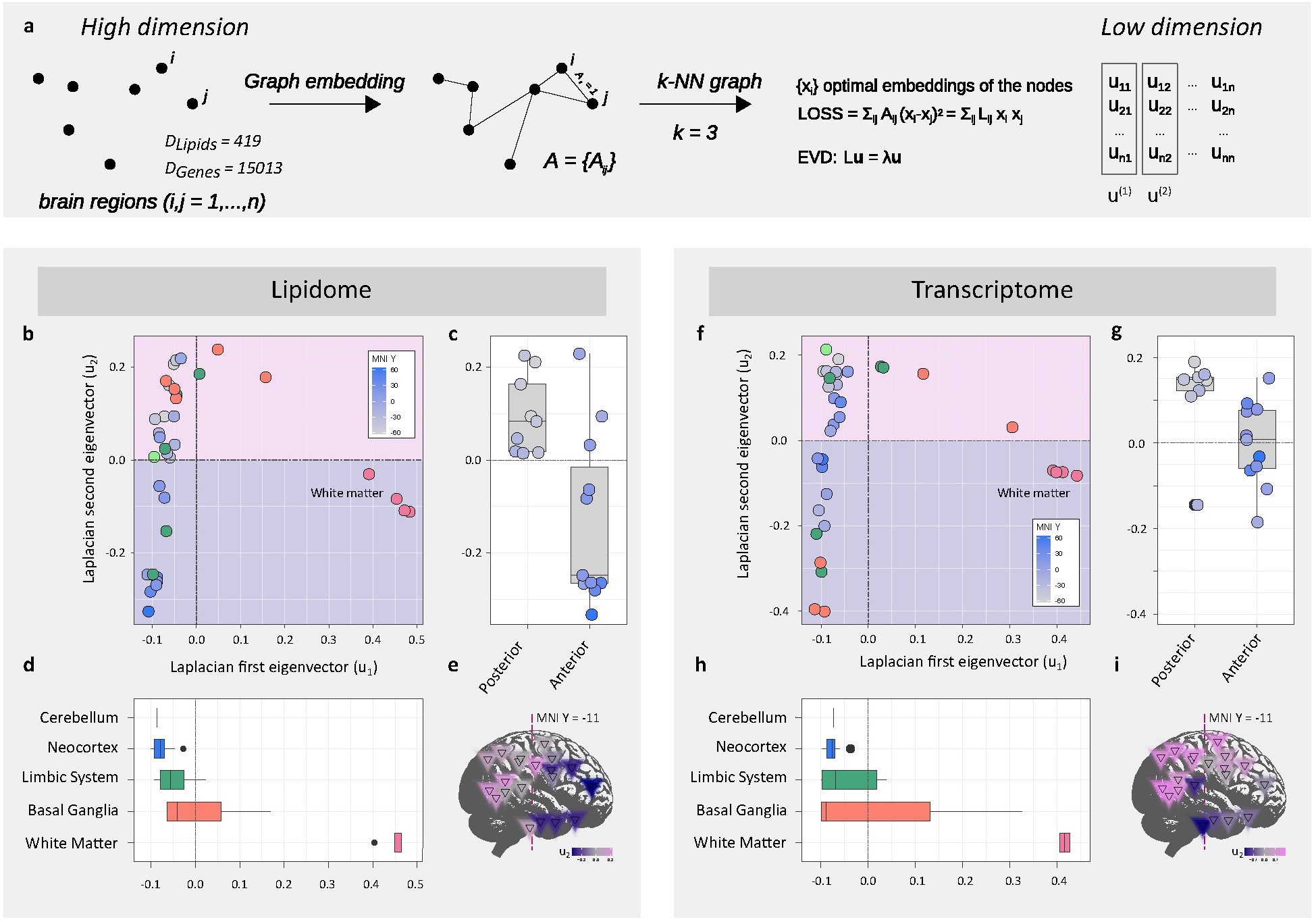
Spectral embedding reveals distinct gradients in lipidomic and transcriptomic maps of brain regions. **a**, Pipeline for spectral embedding: a k-nearest neighbors graph was constructed for brain regions, followed by Laplacian decomposition. The first two non-zero eigenvectors were used as coordinates for embedding. **b**, Spectral embedding of the lipidomic network. Regions separate clearly along two axes, reflecting strong white–gray matter differences and a continuous rostrocaudal gradient in the neocortex. **c-d**, Distributions of lipidome eigenvector (*u*_2_) components plotted along the rostrocaudal axis (c) and across major anatomical structures (d). **e**, Spatial projection of neocortical regions colored by lipidomic eigenvector values, illustrating a smooth anterior–posterior gradient. **f-i**, Equivalent analyses for the transcriptomic network. While broad anatomical separation is preserved, transcriptomic embeddings lack the clear rostrocaudal gradient observed in the lipidome. Together, these analyses show that spectral embeddings capture complementary aspects of brain organization: both lipidome and transcriptome reflect major anatomical divisions, but only the lipidome reveals a continuous cortical gradient.

While the zeroth eigenvector of Laplacian is always constant, the first eigenvector robustly separated white matter from neocortex in both datasets (Figure 2b,f), consistent with the MDS results (Figure 1c,f). As before, two basal ganglia and two limbic regions consistently clustered with white matter. The second eigenvector, however, revealed a clear divergence between modalities. In lipidomic space, limbic regions remained spatially coherent, whereas in transcriptomic space they became maximally separated (Figure 2b,f). This indicates that lipid abundances largely reflect anatomical proximity, whereas transcriptional programs impose functional specialization that overrides physical adjacency.

Within the neocortex, Laplacian embeddings showed another key difference. Neocortical regions formed a vertically aligned cluster, with their separation from white matter captured by the first eigenvector. The second eigenvector encoded a continuous rostrocaudal gradient in the lipidome (negative = anterior, positive = posterior; Figure 2c-e), but only a weak gradient in the transcriptome (Figure 2h-i). This geometric encoding extended to limbic structures, which preserved spatial coherence in lipidomic but not transcriptomic space.

Together, these results show that while both molecular layers preserve the white–gray matter axis, they capture spatial relationships through distinct mechanisms. The lipidome faithfully encodes neuroanatomical geometry, including rostrocaudal gradients and limbic cohesion, whereas the transcriptome is less geometrybound, reflecting functional and possibly cell-type–driven specialization.

### White-gray classification of lipids and transcripts reveals lipid class asymmetry

The robust separation of brain regions into white and gray matter clusters across both lipidomic and transcriptomic networks suggested that this division represents a fundamental organizing principle of brain molecular architecture. To systematically assess how individual molecules align with this axis, we developed a generalized classification framework that projects each lipid or transcript into a two-dimensional space defined by relative abundance in white matter (x-axis) versus neocortex (y-axis) (Figure 3a).

**Figure 3:**
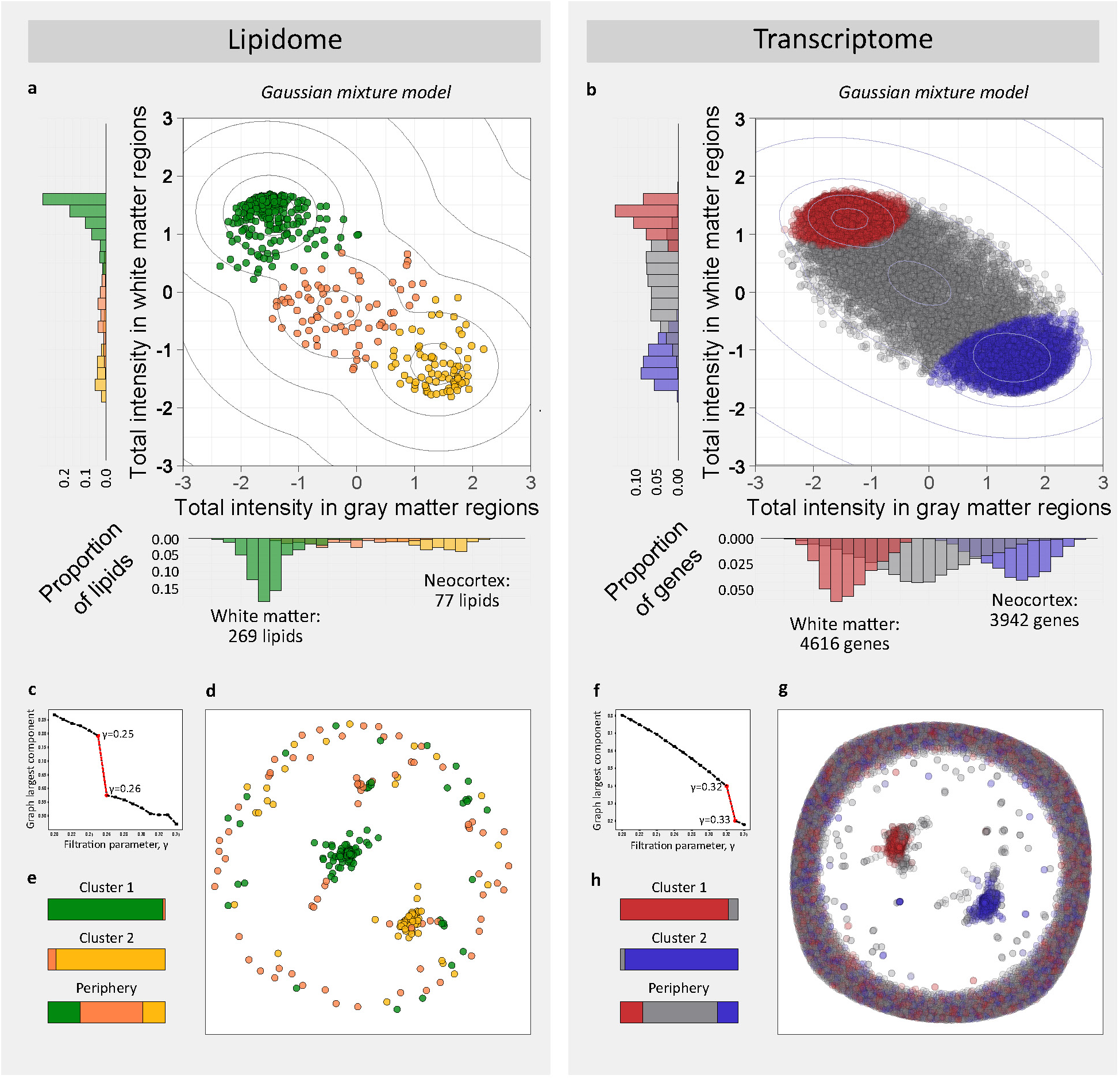
Divergent clustering patterns in lipidomic and transcriptomic networks. **a**, Lipidome embedding based on total signal intensity in gray matter (x-axis) and white matter (y-axis). Gaussian mixture modeling (GMM) identifies three lipid clusters: white matter (green), mixed (orange), and gray matter (yellow). **b**, Analogous embedding of the transcriptome, with GMM clusters defined as white matter (red), mixed (gray), and gray matter (blue). **c**, Percolation analysis of the lipidome network showing a critical threshold (red asterisk) at which a second giant component emerges, used to define optimal filtering. **d**, Lipidome correlation network after filtering for strongest connections. The network separates into two major components and peripheral nodes, colored by GMM clusters. **e**, Mapping of GMM lipid clusters onto the lipidome network structure. **f**, Percolation analysis of the transcriptome network, analogous to c. **g**, Transcriptome correlation network filtered to highlight two major connectivity components and peripheral regions, with nodes colored by GMM clusters. **h**, Mapping of GMM transcriptome clusters onto the transcriptome network. Together, these analyses reveal that while both lipidomic and transcriptomic networks form modular structures, the lipidome is characterized by strong white matter clustering, whereas the transcriptome distributes more evenly across gray and white matter.

Both datasets followed the expected diagonal trend, reflecting overall conservation of white/gray matter composition imposed by data normalization. However, Gaussian mixture modeling (GMM) revealed three distinct molecular classes in each dataset (Figure 3a,b): (i) white matter–enriched, (ii) gray matter–enriched, and (iii) mixed. This analysis uncovered a clear divergence between modalities. The lipidome was strongly biased toward white matter: 65% of lipids (269) were classified as white-enriched, while only 18% (77) were gray-enriched and 17% fell into the mixed category (Figure 3a). This asymmetry reflects the well-documented dominance of lipids in myelinated tracts, where they account for more than 70% of dry tissue mass [73, 74]. By contrast, the transcriptome displayed a more balanced distribution: 30% of genes (4,616) were white-associated, 26% (3,942) gray-associated, and 44% mixed (Figure 3b). These contrasting patterns suggest that while transcripts are more evenly allocated to support diverse functions across regions, lipids remain disproportionately concentrated in white matter to fulfill structural and metabolic roles [33].

To validate the GMM-based classification, we performed an independent graph-based clustering analysis. Molecules were connected by correlation of regional abundance, and networks were progressively filtered until the largest connected component fragmented (Figure 3c,f). At critical thresholds (lipids = 0.26, genes = 0.33), both modalities separated into two large communities. The lipid network showed a pronounced asymmetry: the larger component comprised 241 lipids, 98% of which belonged to the GMM-defined white-enriched cluster, while the smaller contained 58 lipids, 92% aligned with the gray-enriched cluster. The transcriptome network, by contrast, yielded more balanced components: one with 3,023 genes (92% white-associated) and another with 2,347 genes (96% gray-associated). All enrichments were significant by hypergeometric test with Bonferroni correction (*p <* 10*^−^*^3^).

We next tested the robustness of the GMM-based anatomical classifications using leave-one-donor-out (LODO) analyses. In each of the four LODO datasets, GMMs reidentified three clusters for both genes (Figure S6) and lipids (Figure S8) corresponding to white-matter–enriched, gray-matter–enriched, and mixed profiles. Stability, quantified by the Adjusted Rand Index (ARI) against the full-data solution, was high for genes (mean ARI = 0.821; Figure S7) and lipids (mean ARI = 0.924; Figure S9). These results indicate that the three-way partitioning—and the underlying WM/GM asymmetry—is not driven by any single donor.

The strong concordance between the different approaches confirms that the white–gray matter division is a dominant molecular axis in both modalities. Yet, the pronounced asymmetry of the lipidome—with a small but distinct set of 77 gray-enriched lipids—highlights unique lipid specializations in gray matter that warrant further biochemical characterization.

### Decoupling lipidome from transcriptome: Drifters, Anchors and Synchronizers

Having shown that lipidomic and transcriptomic maps share gross structure yet diverge in finer organization—and that these patterns are robust across donors—we next asked how much of regional lipid variation is aligned with gene expression. We frame this as a linear alignment task (not an out-of-sample prediction problem) to quantify lipid–gene coupling on an absolute scale.

For each lipid, we fit a multiple linear regression (MLR) model to its 35-region profile using transcriptome-derived principal components (PCs) as orthogonal regressors (Figure 4b; Methods). To keep the basis compact, we first retained the 150 genes (top 1% by absolute correlation with the lipid) and computed PCs from this subset; this step is strictly an unsupervised dimensionality reduction, not predictive feature selection. Because this filtering used all regions, the resulting *r*^2^ is a descriptive alignment score, not a generalization estimate. To safeguard against spurious associations, we benchmarked *r*^2^ for every lipid against two permutation nulls: a fully randomized shuffle of regional labels and an anatomy-aware shuffle that preserves broad structure of the five anatomical groups and, therefore, regional composition (Methods).

**Figure 4:**
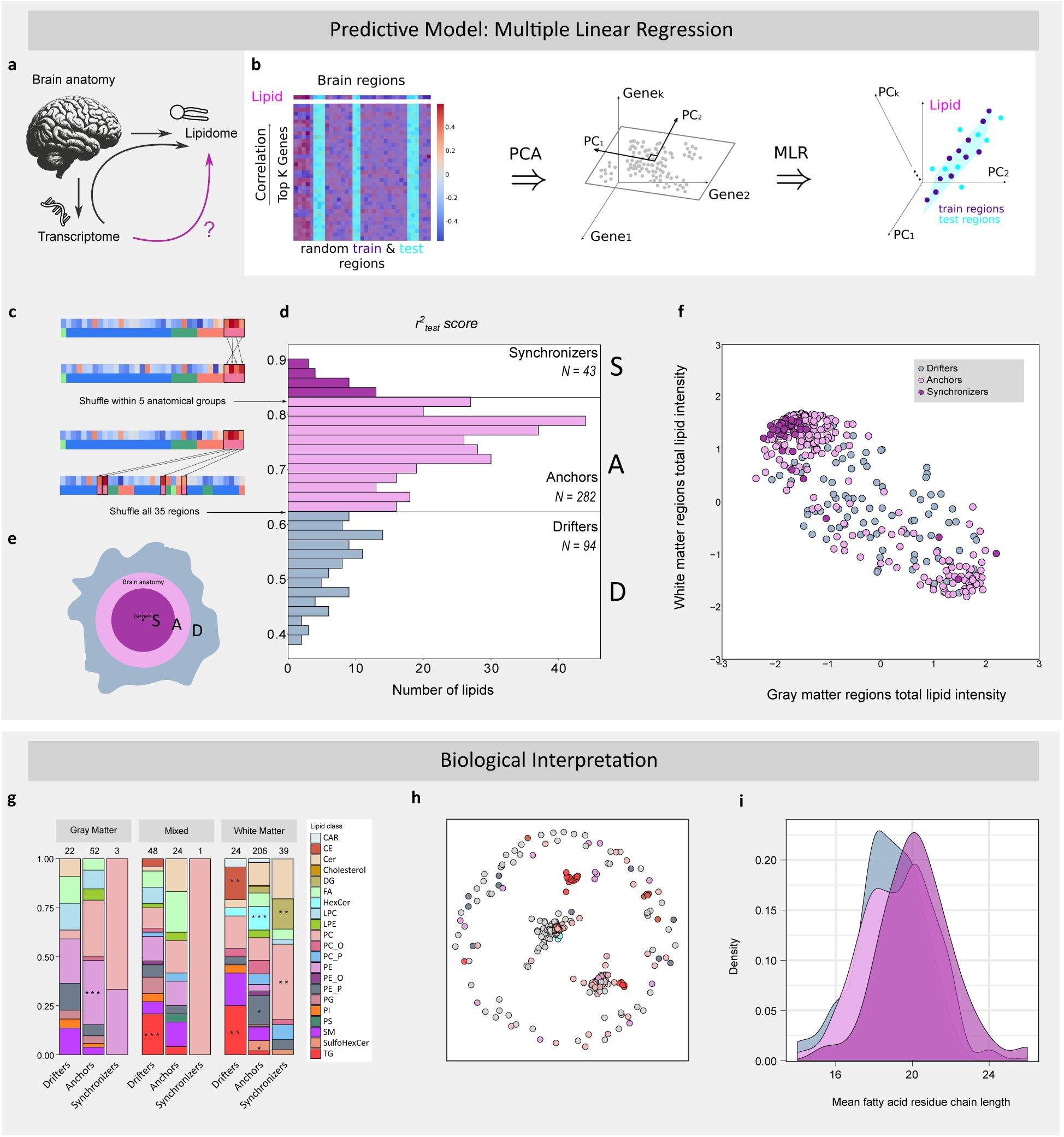
Classification of lipids into Synchronizers, Anchors, and Drifters. **a-b**, Schematic of the MLR pipeline using PCA-reduced gene expression (150 genes; top 1% correlated) to align with lipid levels. Note: the 1% gene filter is an unsupervised dimensionality-reduction step. **c**, Null models based on shuffled lipid concentrations: random shuffling (destroys composition) and anatomy-aware shuffling (preserves regional composition; the Anchor baseline). **d**, Distribution of MLR alignment scores (*r*^2^ scores) defining Synchronizers (gene-coupled), Anchors (anatomy-constrained), and Drifters (independent). **e**, Conceptual diagram of lipid class organization: Synchronizers cluster near transcriptomic programs, Anchors align with broad anatomy, Drifters form a diffuse periphery. **f**, Lipid intensity embedding (axes: neocortex vs. white matter), colored by class. **g**, Enrichment analysis comparing MLR classes and GMM clusters. **h**, Lipid coregulation network colored by bioclass. **i**, Fatty acid chain length distributions for each lipid class. Together, these analyses reveal a tripartite classification of lipids into Synchronizers, Anchors, and Drifters, which are gene-coupled, anatomy/composition-constrained, and transcriptome-independent, respectively.

Across 419 lipids, MLR yielded a unimodal *r*^2^ distribution with a long left tail (Shapiro–Wilk *p <* 10*^−^*^3^; Figure 4d), indicating a subset of lipids poorly predicted by transcriptomic signal. Using the 95th percentiles of the two null distributions as thresholds, we classified lipids into three categories (Figure 4e):

- **Drifters** (23%; *r*^2^ *<* 0.62): lipids showing no significant alignment with gene expression, varying independently of transcriptional programs.
- **Anchors** (67%; 0.62 ≤ *r*^2^ *<* 0.83): lipids predictable at levels consistent with anatomical structure, tethered to regional biology and composition rather than specific genes.
- **Synchronizers** (10%; *r*^2^ ≥ 0.83): lipids strongly coupled to gene expression, exceeding anatomy-based expectations.

Importantly, the anatomy-aware null preserves regional composition (e.g., cell-type mix) within the five anatomical divisions. Thus, Anchors should be interpreted as lipids whose spatial organization is explained by anatomy/composition, whereas Synchronizers exceed this benchmark and Drifters fall below even random expectation.

This framework contrasts with unsupervised clustering, which grouped lipids by lipid–lipid similarity alone. Here, the regression approach reveals the drivers of lipid variation—whether transcriptional programs, anatomical constraints, or neither. Geometrically, Synchronizers form a core tightly linked to transcription, Anchors occupy a surrounding shell constrained by anatomy, and Drifters sit at the periphery, lacking both transcriptomic and lipidomic coherence (Figure 4e). In the following section we demonstrate that this geometry corresponds to latitude on a molecular hypersphere.

The MLR-based classes were robust to donor composition in leave-one-donor-out (LODO) analyses (Figs. S10–S12). Agreement between LODO and full-data labels—quantified by percent intersection and Adjusted Rand Index—was high overall: Anchors and Drifters showed mean intersections of 87% and 85%, respectively, while Synchronizers achieved 52%. The lower stability of Synchronizers reflects their small size and threshold-defined status (exceeding the anatomy-aware null), so reclassifying just a few boundary lipids has a larger proportional effect.

The three classes also showed distinct anatomical distributions (Figure S13). Synchronizers were strongly concentrated in the white matter lipid cluster: 91% of Synchronizers (39 of 43) overlapped with white-enriched lipids (*p <* 10*^−^*^3^). Anchors also favored white matter (73% of 282 Anchors, *p <* 10*^−^*^8^), but with a substantial gray component (18%). By contrast, Drifters overlapped primarily with mixed lipids (51% of 94; *p <* 10*^−^*^3^). In filtered graph embeddings, Drifters localized to the network periphery, consistent with core–shell architectures observed in other complex systems [34, 35] (Figure 4h).

Biochemical enrichment further distinguished the groups (Figure 4g). Anchors were enriched for phosphoethanolamines (PE, *p <* 10*^−^*^3^); Drifters for triacylglycerides (TG, *p <* 10*^−^*^3^); and Synchronizers for phosphatidylcholines (PC, *p <* 10*^−^*^3^) and diacylglycerides (DG, *p <* 10*^−^*^3^). Additional white-matter classes enriched among Anchors included plasmenyl phosphoethanolamines (PE P, *p <* 0.05), sulfohexosylceramides (SulfoHexCer, *p <* 0.05), and hexosylceramides (HexCer, *p <* 10*^−^*^3^). Cholesterol esters (CE, *p <* 10*^−^*^3^) and TG (*p <* 10*^−^*^2^) were specifically enriched among Drifters. All reported enrichments were evaluated by hypergeometric testing with Bonferroni correction (*α* = 0.01).

Chain-length distributions also differed systematically (Figure 4i). Synchronizers peaked at longer fatty acid chains (20 carbons), Drifters at shorter chains (18 carbons), and Anchors displayed a bimodal intermediate pattern (Mann–Whitney U tests, *p <* 0.05). This stratification is consistent with polymer-physics expectations: longer chains experience stronger topological constraints and lower effective mobility, whereas shorter chains reptate more readily and distribute more broadly across regions [41, 37, 36].

In sum, this framework reveals a fundamental decoupling of the lipidome from the transcriptome. Lipids stratify into three mechanistic classes: Synchronizers, tightly coordinated with gene expression; Anchors, constrained by anatomy; and Drifters, uncoupled from either. The enrichment of Synchronizers in myelinated tracts underscores co-regulation of lipid metabolism and gene expression essential for white matter integrity, while Drifters represent a heterogeneous reservoir buffered from transcriptional control.

### A geometric framework for multimodal interactions

Building on the lipid–gene classification, we next examined the structural organization of these interactions in high-dimensional space. After normalization across 35 brain regions, each molecular profile lies on the surface of a 34-dimensional unit sphere (Figure 5a). This projection removes scale differences inherent in cross-regional concentration measurements while preserving biological relationships between molecules. Crucially, the spherical embedding exposes a natural symmetry: two poles corresponding to the white–gray matter axis. Within this geometry, a lipid’s latitude on the hypersphere provides a unifying metric that captures its anatomical associations and degree of transcriptional coupling.

**Figure 5:**
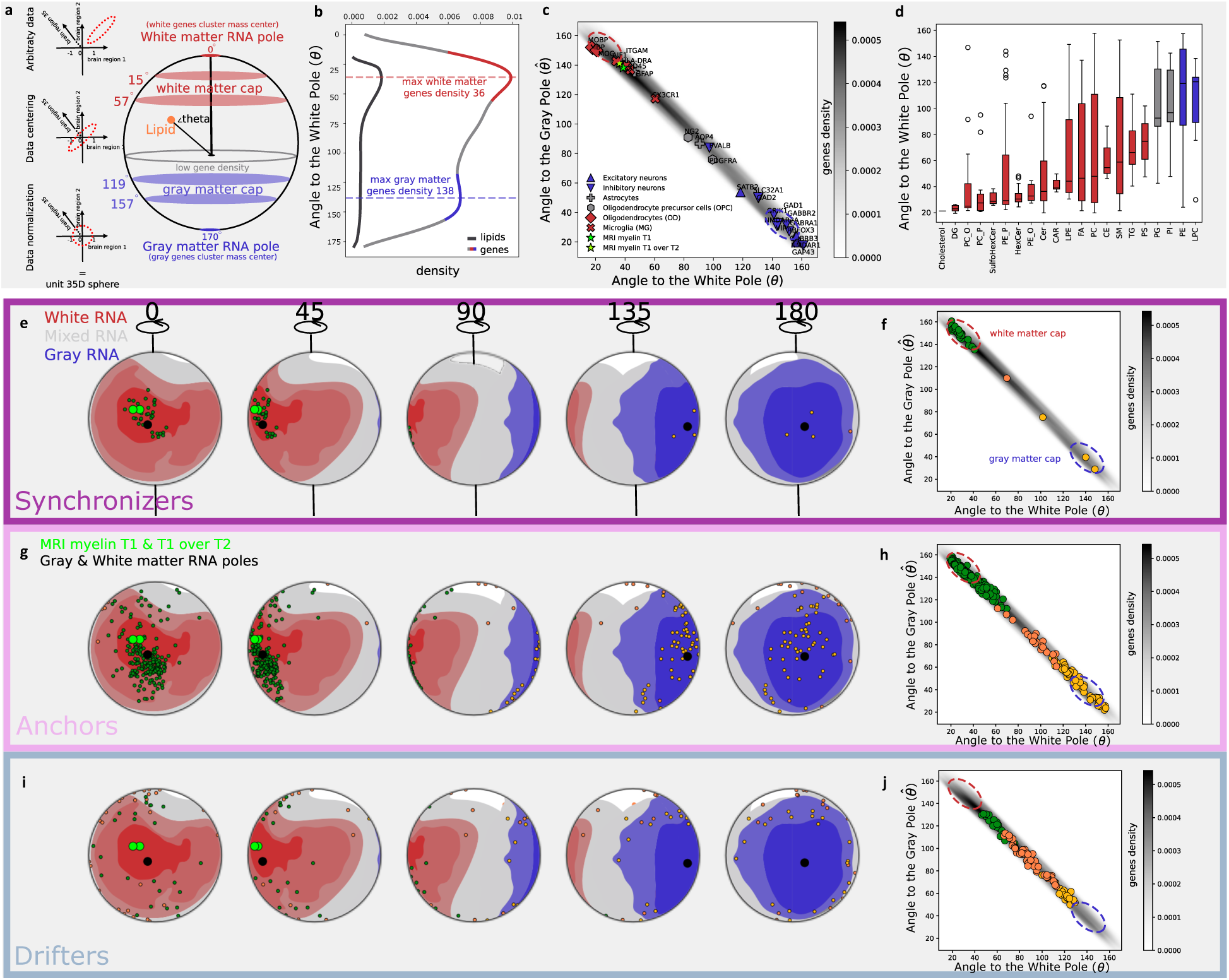
Hyperspherical embedding reveals latent lipid coordinates and category-specific distributions. **a**, Schematic of preprocessing: centering and normalization project molecular vectors onto a unit sphere. Regions of high transcriptomic density are concentrated near the white matter pole (red) and gray matter pole (blue), separated by a low-density equatorial zone (gray). **b**, Gene and lipid density as a function of angular distance to the white matter pole on a 35-dimensional sphere, estimated by spherical kernel density (lipids = black curve; genes = multicolored curve). **c**, Two-dimensional gene probability density map with cell-type marker genes and structural connectome eigenvectors overlaid at their angular coordinates. **d**, Distribution of lipid bioclasses by angular distance to the white matter pole, color-coded by median location: white (red), equatorial (gray), gray matter (blue). **e-f**, Spatial distribution of Synchronizer lipids on the hypersphere, colored by Gaussian mixture model (GMM) class (white = red, gray = blue, mixed = gray). Poles are marked (black dots); background shading reflects transcriptomic density. **g-j**, Equivalent mappings for Anchor and Drifter lipids (as defined by regression-based classification). Together, these analyses show that a single geometric coordinate—angular latitude on the hypersphere — captures the relationship between lipid categories and transcriptomic density, with Synchronizers clustering near poles, Anchors at intermediate latitudes, and Drifters near the equator.

To define coordinates, we set the poles at the centers of mass of the most divergent gene clusters—white matter–enriched and gray matter–enriched genes—separated by ≈170*^◦^*. Each molecular vector’s latitude was measured relative to the white pole, *θ*, with the complementary gray-pole latitude *θ* = *π* − *θ*. We also quantify the nearest-gene angular distance Δ as the geodesic angle to the closest transcript vector. Gene density showed sharp peaks at both poles, forming white and gray matter “caps” separated by an equatorial trough (Figure 5b,f). Lipids, by contrast, were strongly biased toward the white pole, dispersing more gradually toward the gray pole (Figure 5b). This asymmetric distribution mirrors the lipid class imbalance described above (Figure 3), highlighting the link between latitude *θ* and lipid predictability from gene expression.

We next generated a gene density map across latitude pairs (*θ, θ*) (Figure 5c). The diagonal stripe of high density recapitulated the overall gray–white balance, with a slight skew toward the white pole. Cell-type markers aligned along this stripe in expected fashion: neuronal markers clustered near the gray pole, glial markers near the white pole, and astrocyte markers near the equator, consistent with their ubiquitous distribution. Myelin-associated markers displayed particularly distinct patterns, with oligodendrocyte (OD) markers tightly packed at the white pole and oligodendrocyte precursor cells (OPCs) localized equatorially. Neuronal markers spanned ∼50*^◦^*near the gray pole, reflecting subtype heterogeneity. These patterns align closely with established cytoarchitecture.

Projecting lipid bioclasses into the same framework (Figure 5d) revealed a strong white hemisphere bias, dominated by myelin-associated lipids such as cholesterol, diacylglycerols (DG), phosphatidylcholines (PC O, PC P), and sulfatides (SulfoHexCer). Yet some classes diverged: phosphatidylglycerols (PG) and phosphatidylinositols (PI) occupied equatorial positions, suggesting mixed gray–white associations, while phosphatidylethanolamines (PE) and lysophosphatidylcholines (LPC) preferentially localized near the gray pole, consistent with their enrichment in neuronal membranes [38].

Finally, mapping the MLR-based lipid classes onto this hyperspherical framework revealed clear zonation (Figure 5e-i). Synchronizers localized to polar regions of maximal gene density, Anchors occupied intermediate latitudes near high-density zones, and Drifters clustered around the equator, coinciding with minimal gene density. Supporting heatmaps (Figure 5f,h,j) confirmed a systematic pole-to-equator migration across classes, with predictability decreasing in the order Synchronizers → Anchors → Drifters. Thus, angular displacement from the gene white pole, local gene density, and lipid predictability are tightly interdependent—a relationship we validate analytically below.

### Analytical model reveals how hyperspherical geometry governs gene–lipid predictability

The hyperspherical framework provides a geometric explanation for why some lipids are well predicted by gene expression while others are not. Classification into Synchronizers, Anchors, and Drifters reflects each lipid’s position relative to local gene density, which varies systematically with latitude *θ* (Figure 5b). Poorly predictable lipids (Drifters) cluster near the equator (*θ* = *π/*2; Figure 5j), whereas highly predictable Synchronizers concentrate near the poles (*θ* = 0 or *π*; Figure 5f). Anchors occupy intermediate latitudes. The correspondence is not absolute—some Synchronizers appear equatorially and some Anchors near the poles—indicating additional factors modulate predictability.

To capture these relationships more precisely, we explored a second geometric variable: the angular distance Δ to the nearest gene. Geometrically, small Δ values indicate close alignment to gene profiles, and hence higher predictability. By construction, lipids near the poles naturally have smaller Δ, while equatorial lipids generally have larger Δ, although chance proximity to specific genes is possible. Thus, predictability reflects the joint distribution of latitude *θ* and nearest-gene distance Δ.

We formalized this intuition with a probabilistic model of random gene placement on the hypersphere. We modeled an *n* − 1-dimensional sphere (*n* = 35 brain regions) by uniformly sampling angular coordinates (latitude *θ* and *n* − 2 azimuths; Methods). This produces the empirically observed pole-peaked gene density and equatorial depletion. Within this geometry, the expected nearest-neighbor angular distance is:

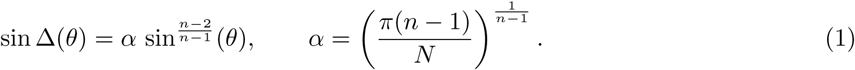

Here *α* is a scaling factor that accounts for finite sample size. With *n* = 35 and *N* = 15, 013, the predicted value is *α* ≈ 0.866.

Remarkably, the theoretical curve predicted by Eq. (1) closely overlapped the overall empirical distribution of lipids (Figure 6). Residuals were near zero for all classes (mean residuals: Synchronizers 0.001 ± 0.04; Anchors 0.005 ± 0.05; Drifters −0.01 ± 0.04), confirming the model’s validity. Thus, the lipid–gene relationships observed in brain data conform closely to expectations from hyperspherical geometry.

**Figure 6:**
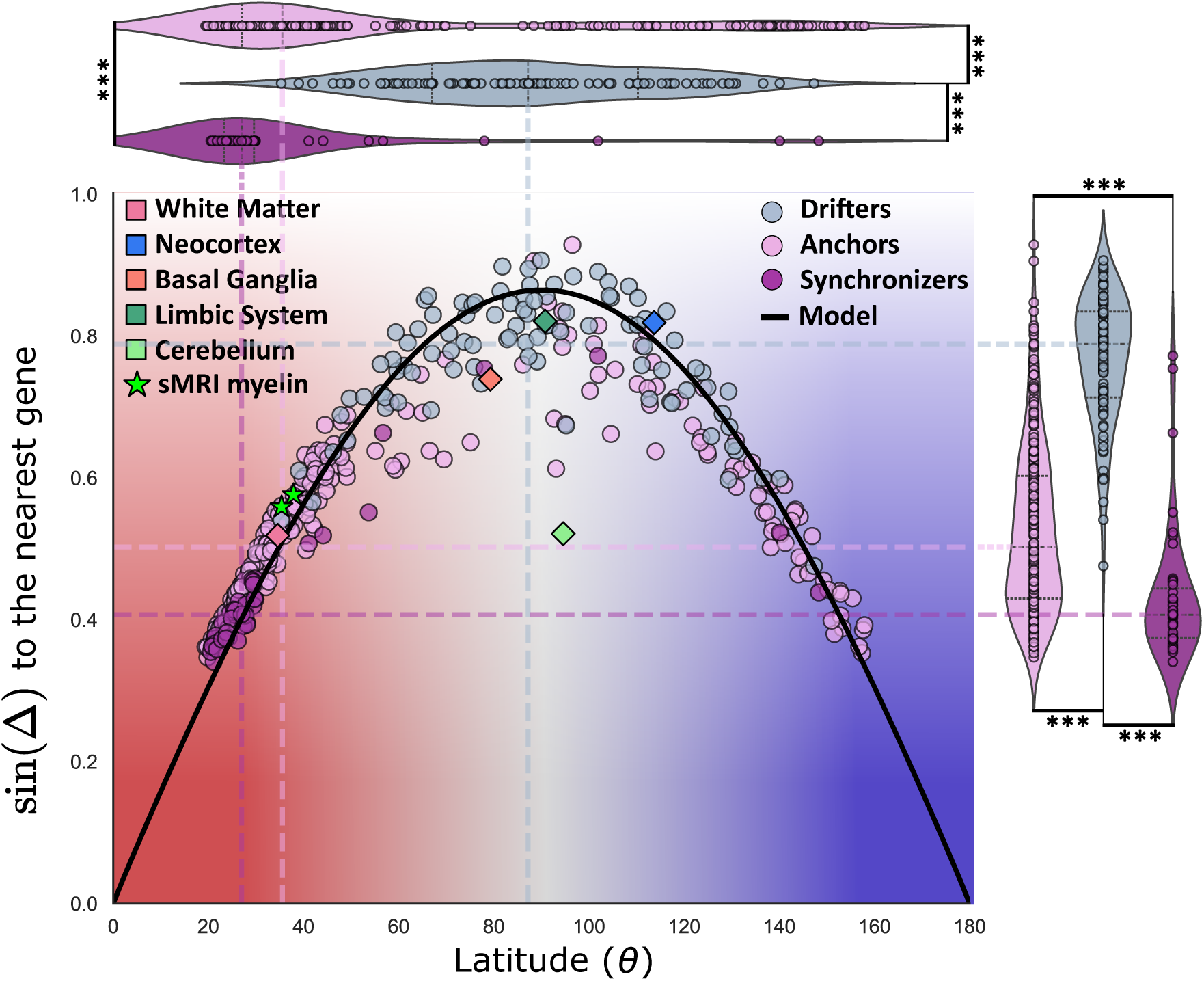
Lipid latitude predicts transcriptomic coupling and aligns with anatomical and imaging features. Scatterplot shows the relationship between lipid latitude on the 34-dimensional hypersphere (x-axis) and average sine-distance to the 150 nearest genes (y-axis). The solid black line indicates model predictions. Background shading reflects transcriptome density zones: red = white matter pole, blue = gray matter pole, white = equatorial region. Green stars denote MRI myelin signals, and colored squares represent anatomical basis vectors (binary encodings of major brain systems). Violin plots (top, right) display latitude and gene-distance distributions for each lipid category (Synchronizers, Anchors, Drifters). Statistical comparisons were made using Mann–Whitney U tests with FDR correction (***, FDR-adjusted *p <* 0.001). Together, these analyses show that lipid predictability increases jointly with decreasing latitude (*θ*) toward the white pole and decreasing nearest-gene distance (Δ), with Synchronizers clustering at low *θ*/Δ, Anchors at intermediate values, and Drifters at high *θ*/Δ. This joint geometric dependence provides a simple interpretable readout of gene coupling, consistent with both imaging and anatomical structure.

We next asked whether lipid classes segregate systematically along the *θ*–Δ curve. Indeed, Synchronizers localized at lower latitudes (median 27*^◦^*) and shorter sin(Δ) (0.40 ± 0.04), Anchors at intermediate positions (36*^◦^*, sin(Δ) = 0.50 ± 0.09), and Drifters near the equator (88*^◦^*, sin(Δ) = 0.81 ± 0.04). Pairwise Mann– Whitney U tests with FDR correction confirmed significant separation between all classes (*p <* 10*^−^*^3^; Figure 6, violin plots). Thus, latitude and local gene density jointly determine lipid predictability.

We then examined deviations from the model by comparing empirical Δ*_e_* and predicted Δ values for each lipid bioclass (Tables S4-S5). Three categories emerged:

1. **Clustering beyond model expectations** (Δ*_e_ <* Δ): Lipid classes showing tighter spatial localization to genes than predicted, mainly equatorial groups such as PC, TG, PS, SM, FA, LPE, and PG. Exceptions included PI and LPC near the gray pole and PE O and CAR near the white pole.
2. **Dispersion beyond model expectations** (Δ*_e_ >* Δ): Classes more spatially diffuse than predicted, largely white pole–associated groups such as DG, PC P, and cholesterol.
3. **Model-conforming distributions** (Δ*_e_* ≈ Δ): Classes whose placement closely matched theoretical expectations, including SulfoHexCer, Cer, HexCer, PE P, PC O near the white pole, and PE near the gray pole.

To test generality, we applied the same framework to basis vectors representing major brain structures (neocortex, basal ganglia, limbic system, white matter, cerebellar cortex). Most conformed to model predictions: white matter (*r*^2^ = 0.95) aligned with the Synchronizer class, while basal ganglia, limbic system, and neocortex (*r*^2^ = 0.82, 0.80, 0.78) matched Anchors. The cerebellar cortex, however, was an outlier: despite equatorial latitude (∼95*^◦^*), it exhibited exceptionally high predictability (*r*^2^ = 0.995), consistent with its unique transcriptomic signature and distinct clustering in earlier analyses (Figure 1).

Finally, we mapped structural connectome eigenvectors from sMRI onto the hypersphere. These vectors localized near the white pole, consistent with their myelin-rich origins, and showed high transcriptomic predictability (*r*^2^ = 0.76–0.79), corresponding to Anchors.

Together, these results show that hyperspherical latitude (*θ*) is the primary correlate of lipid–gene predictability, with nearest-gene distance (Δ) providing an additional local refinement. Lipids near the poles (Synchronizers) synchronize tightly with gene expression, those at intermediate latitudes (Anchors) are constrained by anatomical structure, and equatorial lipids (Drifters) remain largely uncoupled. Exceptions such as the cerebellar cortex highlight how unique molecular enrichment can override geometric expectations. Thus, latitude *θ*—modulated by local gene distance Δ—serves as a interpretable coordinate system for estimating molecular predictability across modalities.

## Discussion

Our findings reveal that the lipidome is not merely a passive reflection of transcriptional programs but represents a partly independent axis of brain organization with its own spatial logic. Whereas transcriptomic maps emphasize developmental lineages and cell-type specificity, lipid distributions introduce additional gradients and clustering hierarchies that cut across these boundaries. The strong white–gray matter asymmetry, continuous rostrocaudal gradient, and limbic-specific clusters demonstrate that lipids encode organizational features absent from gene-expression space. These divergences highlight lipid biology as an additional dimension of brain mapping that complements transcriptomics and proteomics.

Several mechanisms likely underlie this decoupling. First, extensive post-transcriptional and post-translational regulation means that mRNA abundance does not fully dictate downstream molecular states. Proteins, metabolites, and especially lipids are shaped by enzymatic activity, degradation, transport, and feedback loops, such that two regions with similar transcriptomes can maintain distinct lipid profiles. Second, lipid distributions are strongly influenced by tissue physiology and structural constraints—such as myelination, membrane composition, or organelle density—which do not map one-to-one onto gene expression. Together, these mechanisms explain why the lipidome does not simply mirror the transcriptome, but instead encodes an additional organizational layer integrating genetic, cellular, and biophysical determinants.

To connect lipid and gene organization, we classified lipids by how well their regional distributions could be predicted from gene expression. The resulting Synchronizer/Anchor/Drifter taxonomy is defined relative to an anatomy/composition baseline: the anatomy-aware null preserves regional composition, so Anchors are expected to follow anatomy/composition; Synchronizers show transcript-coupling beyond that baseline; Drifters fall below random expectation. Biologically, synchronizers tightly track transcriptomic profiles and include myelin-associated and cell-type–specific lipids that reinforce canonical gene-driven divisions of brain architecture [25, 39, 40]. Drifters, in contrast, are poorly explained by transcriptomic variation, likely reflecting regulation by local physiology or context-dependent processes such as signaling, oxidative stress, or metabolic demand. Anchors represent an intermediate class, partially shaped by gene expression but largely reflecting anatomical context and homeostatic roles. Together, these categories provide a functional vocabulary for human brain lipidome.

Interestingly, we observed that lipids with shorter fatty acid chains are more likely to belong to the Drifter class, whereas Synchronizers tend to be longer molecules. This pattern is consistent with a simple polymer-physics universality: shorter macromolecules experience higher effective mobility than longer chains [36, 37, 41], which may contribute to the weaker spatial coupling of Drifters. This hypothesis warrants experimental testing.

To integrate lipidomic and transcriptomic spaces within a common reference frame, we developed a hyperspherical embedding framework. In this approach, each molecular profile is projected onto the surface of a high-dimensional sphere, with the white–gray matter contrast defining natural poles. Unlike neural-network–based multimodal integration methods [10, 42, 43], this representation is transparent and interpretable: angular distances on the hypersphere directly capture cross-modal relationships without opaque transformations. Within this geometry, a lipid’s latitude (*θ*) provides a global coordinate of predictability, while the nearest-gene angular distance (Δ) refines local coupling. Together, *θ* and Δ recapitulate the Synchronizer–Anchor–Drifter hierarchy, with Synchronizers occupying polar regions of high gene density and small Δ, Anchors at intermediate latitudes with moderate Δ, and Drifters concentrated near the equator with large Δ.

By introducing both a functional classification and a geometric coordinate system, we provide a principled, interpretable framework for multi-omics integration in the brain. Because the representation is simple—profiles are centered and projected onto a shared hypersphere—additional molecular or imaging modalities can be embedded without retraining black-box models, enabling direct tests of how their spatial distributions align with transcriptomic and lipidomic structure. Our interactive portal (Fig. 7) implements this embedding, letting users place lipids, structural-MRI markers, and other readouts on the same coordinates and explore their regional profiles via latitude *θ* and nearest-gene distance Δ. In doing so, the framework doubles as a visualization and hypothesis-generation tool for assessing cross-modal alignment across the brain.

**Figure 7:**
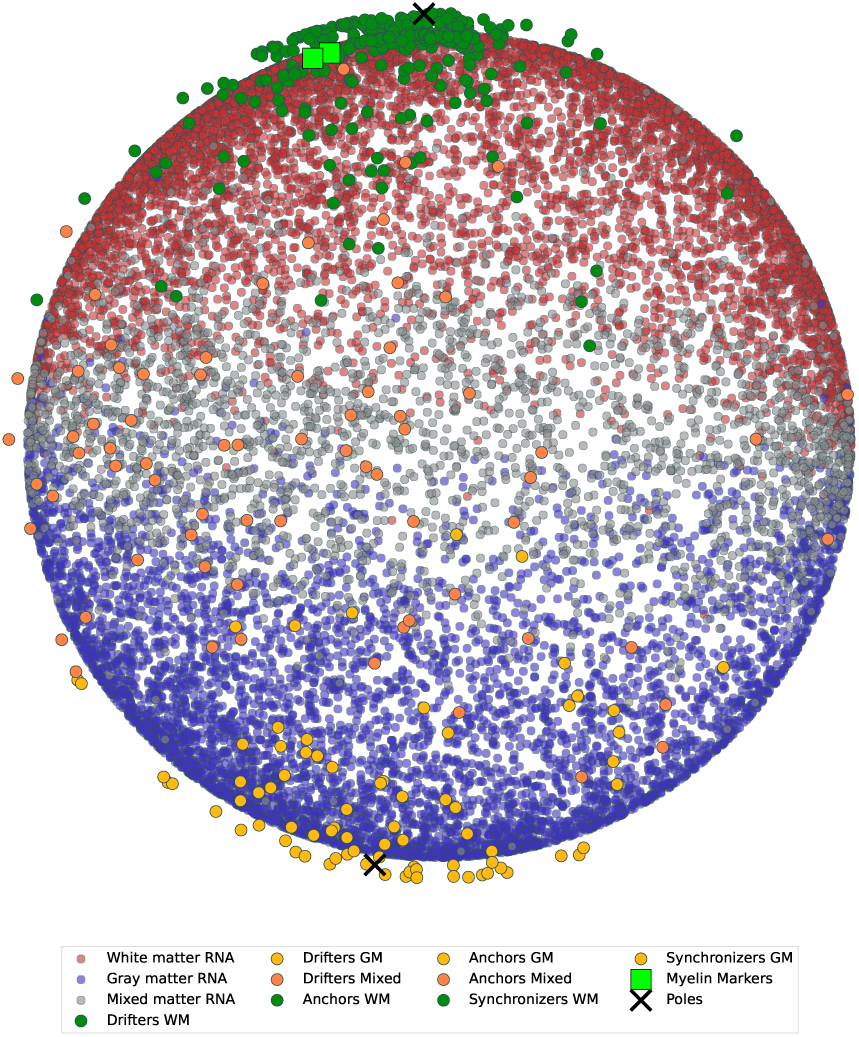
Interactive web portal for 3D molecular mapping. Screenshot of the project’s online platform (https://humanlipidome.pythonanywhere.com), which visualizes transcriptomic and lipidomic data on a shared hyperspherical coordinate system. The portal allows dynamic 3D rotation and zoom, onclick profiling of individual genes or lipids across brain regions, and real-time toggling of molecular layers. This resource enables researchers to explore multimodal relationships directly and to interrogate the spatial organization of lipids and genes in an interactive framework.

Beyond their immediate neuroanatomical implications, these findings open new opportunities for investigating lipid contributions to brain function and disease. Lipidomic alterations are increasingly implicated in neurodegeneration, psychiatric disorders, and aging, yet systematic integration with transcriptomic and imaging data has remained limited. Positioning the lipidome alongside the transcriptome as an equal partner yields a more complete molecular blueprint of the human brain—one that unites genetic programs with biochemical architecture.

## Methods

### Analytical model for the angular distance to the nearest gene

Here we consider a simple probabilistic model of randomly distributed points on a surface of a hypersphere and show that the expected angular distance to the nearest point in the framework of this model recapitulates the experimental data.

In our analytical null, points are sampled uniformly in angular coordinates (not uniformly on surface area), which—through the sin*^n−^*^2^ *θ* surface element—produces pole-enriched density and equatorial depletion. We adopt this on purpose to mirror the empirical gene density; the model is thus a minimal geometric null, not a mechanistic generator.

We consider a unit hypersphere *S^n−^*^1^ in *n*-dimensions, where points are parameterized by *n* − 1 angles: one polar angle *θ* ∈ [0*, π*] and *n*−2 azimuthal angles *ϕ*_1_, · · · *, ϕ_n−_*_2_ ∈ [0, 2*π*]. The infinitesimal surface element on this sphere is as follows:

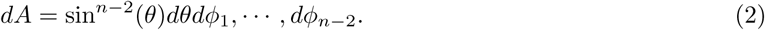

Our probabilistic model assumes that the points are sampled independently and uniformly in the space of the angular parameters *θ* and *ϕ*_1_*, ϕ*_2_, · · · *, ϕ_n−_*_2_. We note that such a model generates uneven distribution of points on the surface of the hypersphere. The reason for that is the positive curvature of the hypersphere and the associated dependence of the infinitesimal surface element on the latitude *θ* (Eq. (2)). Therefore, the actual physical density *ρ*(*θ*) of points in our model scales as:

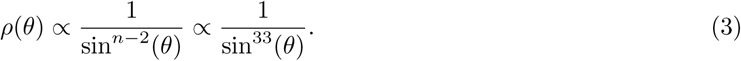

We note that such a simple probabilistic model naturally generates the gene-rich poles (white and gray) as observed in the experimental data. Indeed, for *θ* ≈ 0 and *θ* ≈ *π* the density diverges, reflecting the concentration of the genes at the poles with the opposite behavior. At the equator the density of points is the smallest qualitatively recapitulating the distribution of the genes in the experimental data (5b).

Now we ask if one can describe the typical angular distance Δ to the nearest gene using this model.

Naturally, Δ depends on the latitude *θ* on the hypersphere and, therefore, is dependent on the local density of genes through Eq. (3). For a hypersphere of dimensionality *n*− 1, the area of a spherical cap of the angular size Δ scales as ∝ sin*^n−^*^1^(Δ). Setting the expected number of points in this cap to 1 allows to derive the expected angular distance to the nearest gene at each value of the latitude:

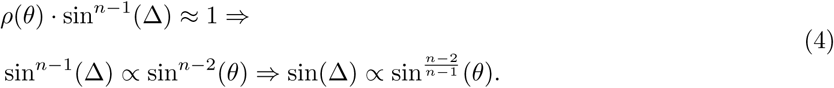

Crucially, for high-dimensional spaces (i.e., large number of brain regions, *n* ≫ 1) the result (4) suggests that the angular distance to the nearest gene Δ is almost perfectly described by the lipid latitude *θ*. For example, in our case *n* = 35, 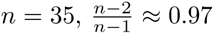 and thus Δ ≈ *θ*. In particular, our model predicts that the sine-distance to the nearest gene scales linearly with sine distance from the lipid to the white pole, sin(Δ) ≈ sin(*θ*).

Figure 6 demonstrates the close agreement between our simple model and the data. The solid black curve represents 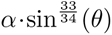 – the expected sine-distance to the nearest gene as predicted by the model. The proportionality constant *α* in this expression can be derived analytically by keeping the prefactors in the argument above. The density of points at latitude *θ* is given by 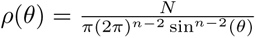, where N is the total number of points. This formula reflects the fact that the uniform sampling of angular coordinates, combined with the surface area element sin*^n−^*^2^(*θ*), leads to a local density inversely proportional to the physical area at each latitude, as explained above. For small angular radii Δ, the area of a cap is approximated by 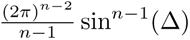. Setting the expected number of points in the cap to unity, we have 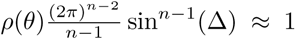, which simplifies to 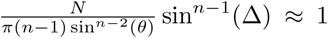. Rearranging yields 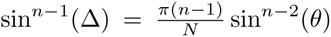 and thus, 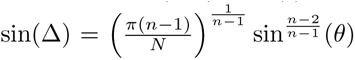. The proportionality constant is therefore 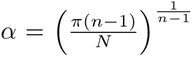. For *n* = 35 and *N* = 15013, this evaluates numerically to *α* ≈ 0.86.

### Multidimensional Scaling

To investigate the structural organization of brain regions across lipidomic and transcriptomic datasets, we applied classical multidimensional scaling (MDS) [44, 45, 46] to pairwise correlation matrices of regional molecular profiles. For each normalized dataset (lipidome and transcriptome), the correlation matrix *C* ∈ *R^n×n^* was computed, where *n* represents the 35 anatomically defined brain regions, and each entry *C_ij_* corresponds to the Pearson correlation coefficient between regions *i* and *j*. Pairwise correlations between brain regions were converted to distances using 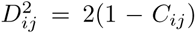, where *C_ij_* is the Pearson correlation coefficient [47]. Classical MDS was then applied to the squared distance matrix, which involves a double-centering procedure to yield a Gram (inner product) matrix amenable to eigendecomposition: 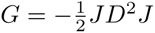, where *J* is a centering matrix. The eigenvectors corresponding to the largest positive eigenvalues of *G* provide coordinates for an optimal embedding in a low-dimensional Euclidean space. The resulting spatial layouts allowed a direct comparison of molecular organization across regions, with axes that captured fundamental partitions such as the white–gray matter divide and finer, region-specific gradations (Figure 1).

### Laplacian Eigenmaps

To further resolve the geometric structure of molecular brain networks, we applied Laplacian Eigenmaps, a nonlinear manifold learning method rooted in spectral graph theory, that preserves both local and global relationships among brain regions. For each molecular dataset, we began by constructing a k-nearest neighbor (k-NN) graph [48], in which each brain region was represented as a node and edges connected each region to its k most molecularly similar neighbors (Figure 2a). Similarity between regional molecular profiles was quantified using pairwise correlation-based distances, ensuring a connectivity pattern that linked regions with similar structural identities, such as those within the neocortex or white matter.

From the adjacency matrix of the k-NN graph, we computed the degree matrix *D* and formed the normalized graph Laplacian *L* = *I* − *D^−^*^1*/*2^*AD^−^*^1*/*2^ [49], where *A* is the adjacency matrix and *I* is the identity matrix. We then solved the eigenvalue problem *Lv* = *λv*, selecting the leading nontrivial eigenvectors (excluding the zeroth constant eigenvector corresponding to *λ*_0_ = 0) to embed the brain regions into a low-dimensional space. These eigenvectors correspond to the smoothest modes of variation on the graph and capture latent spatial gradients present in the molecular data. This spectral embedding enabled us to simultaneously preserve local relationships—such as anatomical community structure – as well as major global gradients like the rostrocaudal (anterior-posterior) axis of the neocortex.

### Gaussian Mixture Model

To systematically classify molecules by their anatomical specificity, we embedded each gene or lipid in a space defined by its relative signal in neocortex versus white matter, capturing their large-scale distribution patterns across the brain. Gaussian Mixture Models (GMMs) were applied to this two-dimensional representation [50, 51, 52]. The GMM algorithm, which models the data as a weighted sum of three Gaussian distributions, probabilistically assigned each molecule to one of three clusters – white matter-enriched, gray matter-enriched, or mixed – revealing the dominant anatomical association of each transcript or lipid species. This framework revealed a striking organizational asymmetry between the two molecular layers, as the lipidome showed a strong white-matter bias (65% of species) compared to the more balanced transcriptome (30% white- vs. 26% gray-enriched).

### Network Filtering Clustering

To further dissect the organizational structure within molecular brain networks, we employed network filtering clustering – a graph-based approach that identifies robust molecular communities through percolation-based thresholding. For each molecular layer, we constructed weighted similarity networks where nodes represented genes or lipids, and edges encoded pairwise Pearson correlations between regional abundance profiles. Networks were iteratively filtered by increasing correlation thresholds, systematically pruning weaker edges until the largest connected component fragmented at a critical percolation threshold. This phase transition (*γ*_lipids_ = 0.26 and *γ*_genes_ = 0.33), marked by abrupt network fragmentation into multiple large components, objectively defined modular boundaries within each molecular network. Hypergeometric testing confirmed significant concordance between network-filtered components and GMM clusters (all p-value *p <* 10*^−^*^10^, Bonferroni-correction *α* = 0.01) [53, 54].

### Principal Component Analysis

To address the high dimensionality and collinearity intrinsic to transcriptome data when predicting lipid abundance, we designed a principal component analysis (PCA)-based pipeline [55, 56, 57, 58]. For computational tractability, the transcriptomic feature space was first filtered for each lipid by retaining the 150 (top 1%) genes exhibiting the strongest regional correlation. Following this initial filtering, Principal Component Analysis (PCA) was applied to the expression matrix of these genes to derive a smaller set of orthogonal, maximally informative components. These components, being statistically orthogonal and maximally informative, served as predictors in subsequent MLR models, improving model stability and interpretability when linking gene expression with lipid variability.

### Multiple Linear Regression

We use multiple linear regression (MLR) to quantify alignment between each lipid’s regional profile and transcriptomic variation across 35 brain regions. We do not aim to build out-of-sample predictors.

Our approach involved three key preprocessing steps to ensure robust model performance: (1) brainwise averaging, where data from four individual brains (unless it is LODO) were aggregated to generate consensus lipid and transcriptome profiles for each brain region; (2) mean-centering, which adjusted each gene and lipid vector to have a mean of zero across regions and (3) *L*_2_ normalization, i.e. scaling each vector to unit variance to mitigate biases arising from differences in measurement scales between genes and lipids. The LODO analysis, confirming that the classification is not predominantly driven by a single donor is done in Figs. S10-S12.

For each lipid, we retain the 150 genes (top 1%) with the largest absolute regional correlation to the lipid, then compute principal components (PCs) on this subset. This filtering and PCA are unsupervised dimensionality-reduction steps that provide an orthogonal basis; they are not predictive feature selection. These PCs served as orthogonal predictors *x* in subsequent regression modeling of lipids *y*. Then the MLR model is as follows 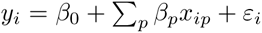, where *β*_0_ is the intercept, *β_p_* are regression coefficients, and *ε_i_* represents residual errors.

To evaluate model performance, we employed a repeated random sub-sampling protocol. This process was iterated 10,000 times, with the 35 brain regions randomly partitioned into a training set (28 regions; ∼ 80%) and a test set (7 regions; ∼ 20%) in each run. The final coefficient of determination (*r*^2^), computed on the test set and averaged across all 10,000 splits, was interpreted as a descriptive measure of lipid–gene alignment rather than a metric of out-of-sample generalization [63]. To assess significance and guard against spurious associations, we compared these observed *r*^2^ values to two permutation-based nulls (random and anatomy-aware; see below) and used their 95th percentiles (*r*^2^ = 0.62 and *r*^2^ = 0.83) as the thresholds for class assignment.

To establish significance, we pass two permutation nulls through the identical pipeline (including the 1% filter and PCA): (i) a fully randomized shuffle of regional labels and (ii) an anatomy-aware shuffle preserving composition within the five anatomical divisions. The 95th percentiles of these null distributions define Drifters (*r*^2^ *<* 0.62; below random), Anchors (0.62 *< r*^2^ *<* 0.83; between random and anatomy-aware), and Synchronizers (*r*^2^ *>* 0.83; above anatomy-aware).

## Author Contributions

K.P. and P.K. designed the research, M.O., A.O., E.S. and K.P. analyzed the data. M.O., A.O., P.K. and K.P. contributed to writing the paper.

## Data Availability Statement

The data analyzed in this study are available at https://zenodo.org/records/10908108

## Code Availability

A detailed step-by-step record of the analysis performed in this study is available at https://github.com/ArseniiOnuchin/human_lipidome

## References

[1] Barabási, D. L., et al. Neuroscience needs network science. J. Neurosci. 43, 5989–5995 (2023).

[2] Cui, H. et al. Towards multimodal foundation models in molecular cell biology. Nature 640, 623–633 (2025).

[3] Barabási, A.-L., Gulbahce, N. & Loscalzo, J. Network medicine: a network-based approach to human disease. Nat. Rev. Genet. 12, 56–68 (2011).

[4] Oltvai, Z. N. & Barabási, A.-L. Systems biology. Life’s complexity pyramid. Science 298, 763–764 (2002).

[5] Barabási, A.-L. & Oltvai, Z. N. Network biology: understanding the cell’s functional organization. Nat. Rev. Genet. 5, 101–113 (2004).

[6] Ravasz, E., Somera, A. L., Mongru, D. A., Oltvai, Z. N. & Barabasi, A.-L. Hierarchical organization of modularity in metabolic networks. arXiv [cond-mat.soft] (2002).

[7] Cho, K. et al. Network analysis of the metabolome and transcriptome reveals novel regulation of potato pigmentation. J. Exp. Bot. 67, 1519–1533 (2016).

[8] Zhang, S. et al. Network analysis of metabolome and transcriptome revealed regulation of different nitrogen concentrations on hybrid poplar cambium development. Int. J. Mol. Sci. 25, 1017 (2024).

[9] Schaffer, L. V. et al. Multimodal cell maps as a foundation for structural and functional genomics. Nature 642, 222–231 (2025).

[10] Tang, X. et al. Explainable multi-task learning for multi-modality biological data analysis. Nat. Commun. 14, 2546 (2023).

[11] Seidlitz, J. et al. Transcriptomic and cellular decoding of regional brain vulnerability to neurogenetic disorders. Nat. Commun. 11, 3358 (2020).

[12] Arkhipov, A. et al. Integrating multimodal data to understand cortical circuit architecture and function. Nat. Neurosci. 28, 717–730 (2025).

[13] Chen, J. et al. Integration of multimodal data for deciphering brain disorders. Annu. Rev. Biomed. Data Sci. 4, 43–56 (2021).

[14] Horgusluoglu, E. et al. Integrative metabolomics-genomics approach reveals key metabolic pathways and regulators of Alzheimer’s disease. Alzheimers. Dement. 18, 1260–1278 (2022).

[15] Gabitto, M. I. et al. Integrated multimodal cell atlas of Alzheimer’s disease. Nat. Neurosci. 27, 2366–2383 (2024).

[16] Chierici, M. et al. Integrative Network Fusion: A multi-omics approach in molecular profiling. Front. Oncol. 10, 1065 (2020).

[17] Polovnikov, K., Gorsky, A., Nechaev, S., Razin, S. V. & Ulianov, S. V. Non-backtracking walks reveal compartments in sparse chromatin interaction networks. Sci. Rep. 10, 11398 (2020).

[18] Onuchin, A. A., Chernizova, A. V., Lebedev, M. A. & Polovnikov, K. E. Communities in C. elegans connectome through the prism of non-backtracking walks. Sci. Rep. 13, 22923 (2023).

[19] Nechaev, S. K. & Polovnikov, K. Rare-event statistics and modular invariance. Phys.–Usp. 61, 99–104 (2018).

[20] Wang, M., Wang, H. & Zheng, H. A mini review of node centrality metrics in biological networks. International Journal of Network Dynamics and Intelligence 1, 99–110 (2022).

[21] Bertolero, M. A., Yeo, B. T. T. & D’Esposito, M. The modular and integrative functional architecture of the human brain. Proc. Natl. Acad. Sci. U. S. A. 112, E6798–807 (2015).

[22] Kovács, I. A., et al. Network-based prediction of protein interactions. Nat. Commun. 10, 1240 (2019).

[23] Hawrylycz, M. et al. Canonical genetic signatures of the adult human brain. Nat. Neurosci. 18, 1832–1844 (2015).

[24] Roy, M. et al. Proteomic analysis of postsynaptic proteins in regions of the human neocortex. Nat. Neurosci. 21, 130–138 (2018).

[25] Poitelon, Y., Kopec, A. M. & Belin, S. Myelin fat facts: An overview of lipids and fatty acid metabolism. Cells 9, 812 (2020).

[26] Bozek, K. et al. Organization and evolution of brain lipidome revealed by large-scale analysis of human, chimpanzee, macaque, and mouse tissues. Neuron 85, 695–702 (2015).

[27] Huisman, S. M. H. et al. BrainScope: interactive visual exploration of the spatial and temporal human brain transcriptome. Nucleic Acids Res. 45, e83 (2017).

[28] Hawrylycz, M. J. et al. An anatomically comprehensive atlas of the adult human brain transcriptome. Nature 489, 391–399 (2012).

[29] Bernard, A. et al. Transcriptional architecture of the primate neocortex. Neuron 73, 1083–1099 (2012).

[30] Krzakala, F. et al. Spectral redemption in clustering sparse networks. Proc. Natl. Acad. Sci. U. S. A. 110, 20935–20940 (2013).

[31] Saade, A., Krzakala, F. & Zdeborová, L. Spectral density of the non-backtracking operator on random graphs. EPL 107, 50005 (2014).

[32] Vinh, N. X., Epps, J., & Bailey, J. Information Theoretic Measures for Clusterings Comparison: Variants, Properties, Normalization and Correction for Chance. J. Mach. Learn. Res. 11, 2837–2854 (2010).

[33] Osetrova, M. et al. Mass spectrometry imaging of two neocortical areas reveals the histological selectivity of schizophrenia-associated lipid alterations. Consort. Psychiatr. 5, 4–16 (2024).

[34] Polovnikov, K., Kazakov, V., & Syntulsky, S. Core–periphery organization of the cryptocurrency market inferred by the modularity operator. Physica A: Statistical Mechanics and its Applications 540, 123075 (2020).

[35] Borgatti, S. P., & Everett, M. G. Models of core/periphery structures. Social networks 21, 375–395 (2000).

[36] Tamm, M. V. & Polovnikov, K. Dynamics of polymers: Classic results and recent developments. In Order, Disorder and Criticality 113–172 (WORLD SCIENTIFIC, 2018).

[37] Khokhlov, A. R., Grosberg, A. Y., & Pande, V. S. Statistical physics of macromolecules (Vol. 1). New York: AIP press (1994).

[38] Strakova, J., Demizieux, L., Campenot, R. B., Vance, D. E. & Vance, J. E. Involvement of CTP:phosphocholine cytidylyltransferase-beta2 in axonal phosphatidylcholine synthesis and branching of neurons. Biochim. Biophys. Acta 1811, 617–625 (2011).

[39] Dimas, P. et al. CNS myelination and remyelination depend on fatty acid synthesis by oligodendrocytes. Elife 8, (2019).

[40] Barnes-Vélez, J. A., Aksoy Yasar, F. B. & Hu, J. Myelin lipid metabolism and its role in myelination and myelin maintenance. Innovation (Camb.) 4, 100360 (2023).

[41] Grosberg, A. Y., Nechaev, S. K. & Shakhnovich, E. I. The role of topological constraints in the kinetics of collapse of macromolecules. Journal Phys. 49, 2095–2100 (1988).

[42] Ashuach, T. et al. MultiVI: deep generative model for the integration of multimodal data. Nat. Methods 20, 1222–1231 (2023).

[43] Esser-Skala, W. & Fortelny, N. Reliable interpretability of biology-inspired deep neural networks. NPJ Syst. Biol. Appl. 9, 50 (2023).

[44] Torgerson, W. S. Theory and Methods of Scaling. (John Wiley & Sons, Nashville, TN, 1958).

[45] Mardia, K.V. Rigorous Treatment of Geometric Properties (critical for Centering and Eigenvalues). Biometrika 53, 3–4 (1966).

[46] Kruskal, J. B. & Wish, M. Multidimensional Scaling. (SAGE Publications, Thousand Oaks, CA, 1978).

[47] De Leeuw, J. Applications of Convex Analysis to Multidimensional Scaling. (1977).

[48] Connor, M. & Kumar, P. Fast construction of k-nearest neighbor graphs for point clouds. IEEE Trans. Vis. Comput. Graph. 16, 599–608 (2010).

[49] Belkin, M. & Niyogi, P. Laplacian eigenmaps for dimensionality reduction and data representation. Neural Comput. 15, 1373–1396 (2003).

[50] Liu, C. Maximum likelihood estimation from incomplete data via em-type algorithms. in Advanced Medical Statistics 1051–1071 (WORLD SCIENTIFIC, 2003).

[51] Fraley, C. & Raftery, A. E. Model-based clustering, discriminant analysis, and density estimation. J. Am. Stat. Assoc. 97, 611–631 (2002).

[52] Aranda, S., Linares, J.-M. & Sprauel, J.-M. Likelihood maximization against the probability density function shape. in Series on Advances in Mathematics for Applied Sciences 7–13 (WORLD SCIENTIFIC, 2009).

[53] Radicchi, F. Predicting percolation thresholds in networks. Phys. Rev. E Stat. Nonlin. Soft Matter Phys. 91, 010801 (2015).

[54] Dickey, J. M. Multiple hypergeometric functions: Probabilistic interpretations and statistical uses. J. Am. Stat. Assoc. 78, 628 (1983).

[55] Pearson, K. LIII. On lines and planes of closest fit to systems of points in space. Lond. Edinb. Dublin Philos. Mag. J. Sci. 2, 559–572 (1901).

[56] Hotelling, H. Analysis of a complex of statistical variables into principal components. J. Educ. Psychol. 24, 417–441 (1933).

[57] Jolliffe, I. T. Discarding variables in a principal component analysis. I: Artificial data. J. R. Stat. Soc. Ser. C. Appl. Stat. 21, 160 (1972).

[58] Jolliffe, I. T. Discarding Variables in a Principal Component Analysis. II: Real Data. J. R. Stat. Soc. Ser. C. Appl. Stat. 22, 21 (1973).

[59] Krzywinski, M. & Altman, N. Multiple linear regression: Points of Significance. Nat. Methods 12, 1103–1104 (2015).

[60] Yan, X. & Su, X. G. Linear Regression Analysis: Theory and Computing. (World Scientific Publishing, Singapore, Singapore, 2009).

[61] Jolliffe, I. T. A note on the use of principal components in regression. J. R. Stat. Soc. Ser. C. Appl. Stat. 31, 300 (1982).

[62] Xu, Q.-S. & Liang, Y.-Z. Monte Carlo cross validation. Chemometr. Intell. Lab. Syst. 56, 1–11 (2001).

[63] Nagelkerke, N. J. D. A note on a general definition of the coefficient of determination. Biometrika 78, 691 (1991).

[64] Millar, T. et al. Tissue and organ donation for research in forensic pathology: the MRC Sudden Death Brain and Tissue Bank. J. Pathol. 213, 369–375 (2007).

[65] Wang, Q. et al. Ex vivo instability of lipids in whole blood: preanalytical recommendations for clinical lipidomics studies. J. Lipid Res. 64, 100378 (2023).

[66] Caballero-Moreno, L., Luna, A. & Legaz, I. Lipidomes in cadaveric decomposition and determination of the postmortem interval: A systematic review. Int. J. Mol. Sci. 25, (2024).

[67] Mai, J. K. & Majtanik, M. Myeloarchitectonic maps of the human cerebral cortex registered to surface and sections of a standard atlas brain. Transl. Neurosci. 14, 20220325 (2023).

[68] Mai, J. K., Majtanik, M. & Paxinos, G. Atlas of the Human Brain. (Academic Press, San Diego, CA, 2015).

[69] Smith, C. A., Want, E. J., O’Maille, G., Abagyan, R. & Siuzdak, G. XCMS: Processing mass spectrometry data for metabolite profiling using nonlinear peak alignment, matching, and identification. Anal. Chem. 78, 779–787 (2006).

[70] Züllig, T., Trötzmüller, M. & Köfeler, H. C. Lipidomics from sample preparation to data analysis: a primer. Anal. Bioanal. Chem. 412, 2191–2209 (2020).

[71] Khrameeva, E. et al. Single-cell-resolution transcriptome map of human, chimpanzee, bonobo, and macaque brains. Genome Res. 30, 776–789 (2020).

[72] Shimada, M., Omae, Y., Kakita, A., Gabdulkhaev, R., Hitomi, Y., Miyagawa, T., Honda, M., Fujimoto, A., & Tokunaga, K. Identification of region-specific gene isoforms in the human brain using long-read transcriptome sequencing. Science Advances 10, eadj5279 (2024).

[73] Chrast, R., Saher, G., Nave, K. A., & Verheijen, M. H. G. Lipid metabolism in myelinating glial cells: lessons from human inherited disorders and mouse models. Journal of Lipid Research 52, 419–434 (2011).

[74] Duncan, I. D. Inherited and acquired disorders of myelin: The underlying myelin biology. Experimental Neurology 283, 496–505 (2016).

