## Supplementary Materials for "Hyperspherical geometry positions the lipidome as a partly independent axis of human brain organization"

### Supplementary Information for "Hyperspherical geometry positions the lipidome as a partly independent axis of human brain organization"

Maria Osetrova, Arsenii Onuchin, Elena Stekolshchikova, Philipp Khaitovich, Kirill Polovnikov

### Supplementary Figures

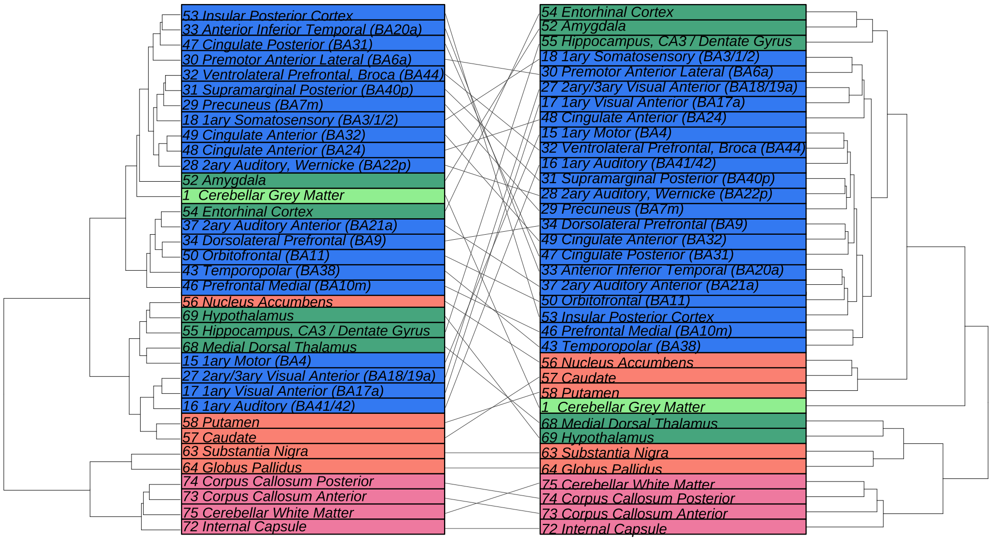


Supplementary Figure 1: Comparative Hierarchical Clustering of Human Brain Regions. This figure presents detailed dendrograms illustrating the hierarchical clustering of 35 human brain regions based on lipidomic (left panel) and transcriptomic (right panel) similarity. The branching structure of the dendrograms reflects the degree of molecular similarity between regions, with shorter branches indicating higher similarity. Brain regions are color-coded according to their major anatomical division: neocortex (royal blue), limbic system (dark sea green), basal ganglia (red-orange), white matter (pink), and cerebellar gray matter (light green). This is an expanded view of the summary diagram in Figure 1d, providing the specific names for each brain region to allow for detailed comparison of how the lipidome and transcriptome group these structures. While both lipidomic and transcriptomic profiles broadly align with the major anatomical divisions of the brain, this detailed comparison reveals subtle but significant differences. For example, the cerebellar cortex forms a distinct outgroup in the transcriptomic tree but not in the lipidomic one. Similarly, limbic regions like the medial dorsal thalamus and hypothalamus cluster with white matter in the transcriptomic network, but not in the lipidomic network. These divergences indicate that the lipidome is not merely a reflection of the transcriptome but constitutes a complementary organizational layer, capturing structural and functional relationships that are not apparent from gene expression alone.

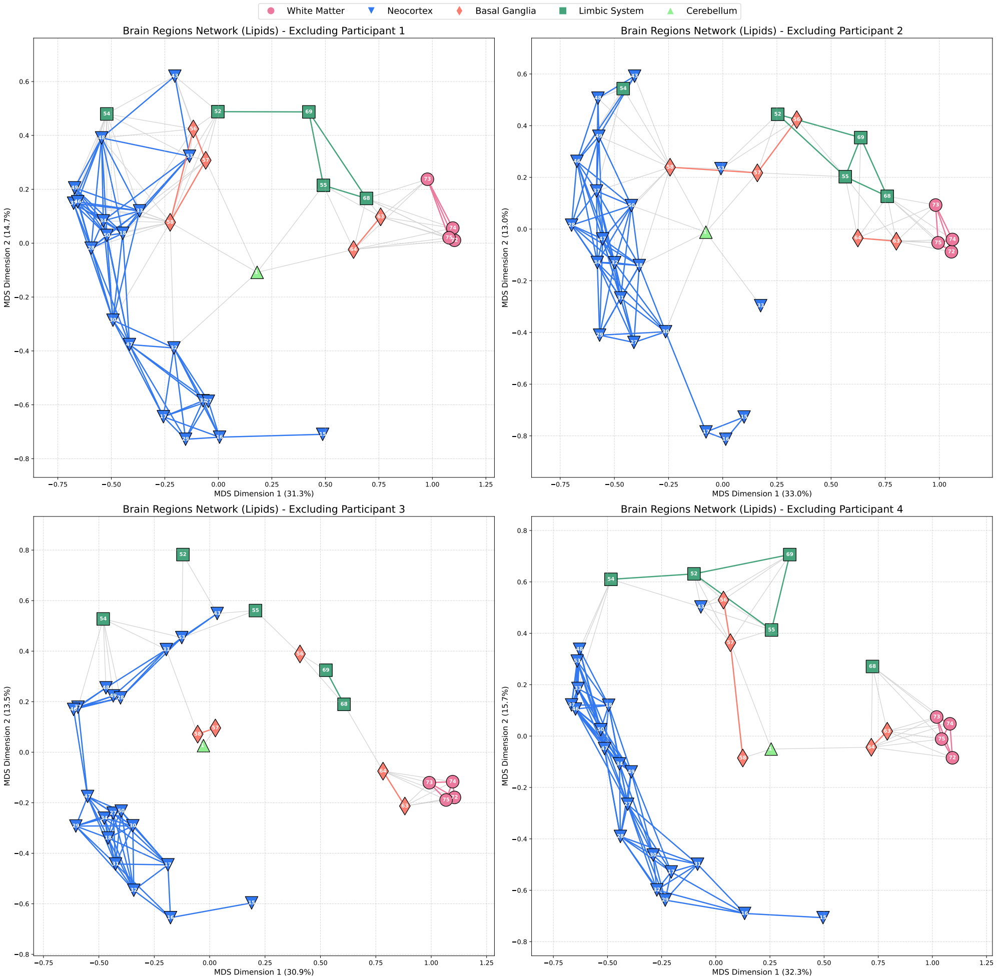


Supplementary Figure 2: Stability of the Human Brain Lipidomic Network Assessed by Leave-One-Out Cross-Validation. This figure displays four network maps illustrating the molecular similarity between 35 human brain regions based on their lipidomic profiles, with each panel representing an experiment where one of four different participants was excluded from the analysis. Node positions are determined by classical Multidimensional Scaling (MDS), such that the distance between nodes reflects their lipidomic dissimilarity. Edges are drawn between regions that are closer than a minimal connection threshold (so network still connected), highlighting the most robust molecular relationships. Nodes are color-coded and shaped by their major anatomical division: neocortex (royal blue, triangle), limbic system (dark sea green, square), basal ganglia (red-orange, diamond), white matter (pink, circle), and cerebellum (light green, inverted triangle). The axes represent the first two MDS dimensions, with the percentage of total variance explained by each dimension noted. The consistency across all four leave-one-out experiments demonstrates the stability of the brain's lipidomic organization. Key anatomical structures, such as the clear separation of white and gray matter and the clustering of neocortical regions, are preserved even when individual datasets are removed.

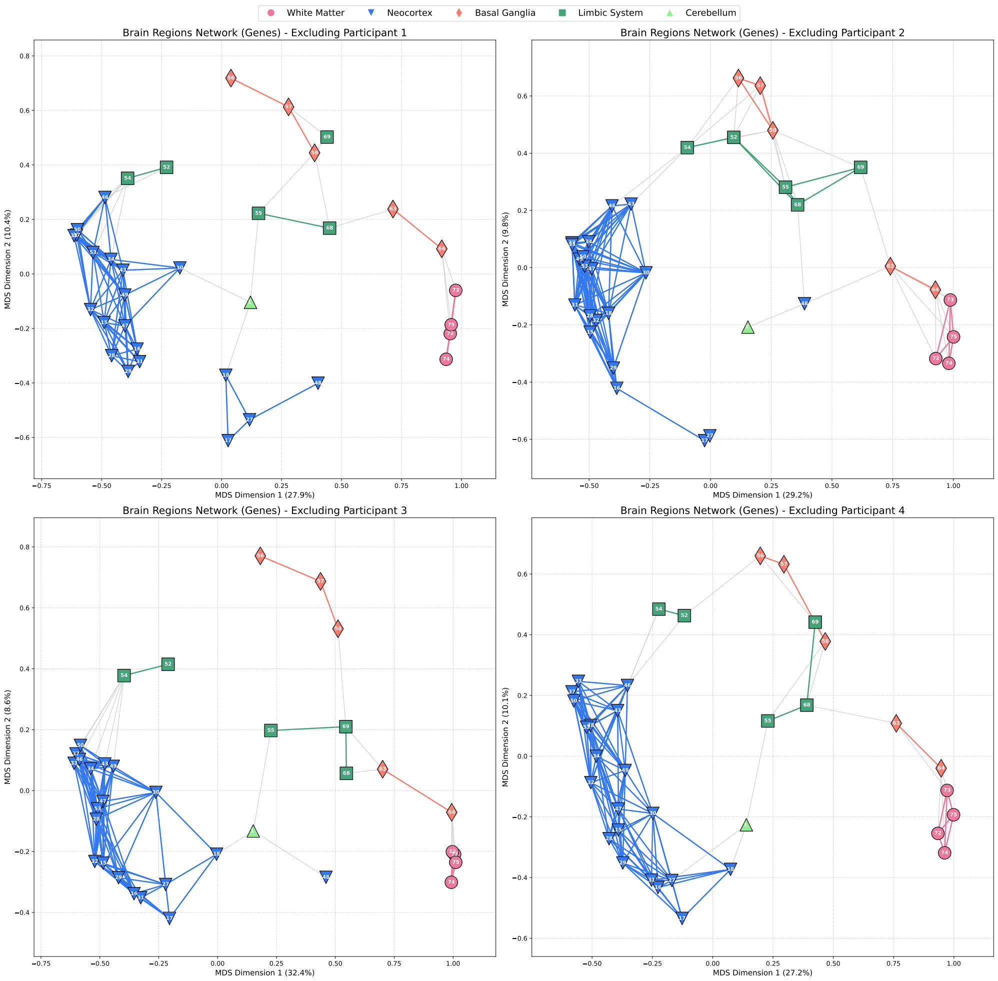


Supplementary Figure 3: Stability of the Human Brain Transcriptomic Network Assessed by Leave-One-Out Cross-Validation. This figure displays four network maps illustrating the molecular similarity between 35 human brain regions based on their transcriptomic profiles, with each panel representing an experiment where one of four different participants was excluded from the analysis. To facilitate direct visual comparison with the lipidomic network (Figure 2), the horizontal axis of each plot has been mirrored. Node positions are determined by classical Multidimensional Scaling (MDS), with the distance between nodes reflecting their dissimilarity in gene expression. Edges are drawn between regions that are closer than a minimal connection threshold, highlighting the most robust transcriptomic relationships. Nodes are color-coded and shaped by their major anatomical division: neocortex (royal blue, triangle), limbic system (dark sea green, square), basal ganglia (red-orange, diamond), white matter (pink, circle), and cerebellum (light green, inverted triangle). The structural consistency across all four leave-one-out experiments confirms that the brain's transcriptomic organization is a stable and reproducible biological feature.

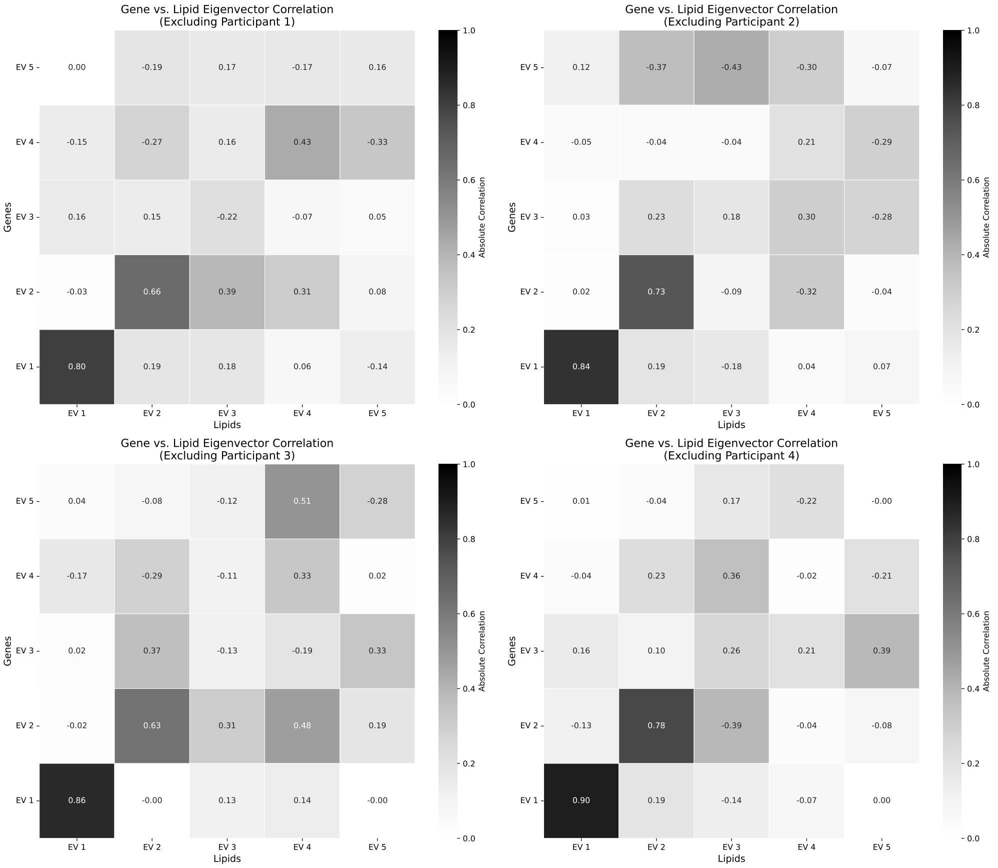


Supplementary Figure 4: Cross-Modal Alignment of Transcriptomic and Lipidomic Principal Components. This figure presents a set of four 5x5 heatmaps, each illustrating the pairwise correlation between the top five eigenvectors derived from the transcriptomic and lipidomic profiles of the human brain. Each panel corresponds to a leave-one-out cross-validation experimen. The components were obtained via classical Multidimensional Scaling (MDS) on the inter-regional distance matrices for genes and lipids, respectively. The color intensity of each cell represents the absolute correlation coefficient, while the annotated number provides the original signed value. This analysis reveals a remarkably strong and stable alignment between the principal axes of variation in the brain's transcriptome and lipidome. Specifically, the first principal component (EV1) of the transcriptome consistently shows a very high positive correlation with the first component of the lipidome across all four experiments (r = 0.80, 0.84, 0.86, and 0.90). A robust, secondary alignment is also evident between the second principal components (EV2), with correlation coefficients of 0.66, 0.73, 0.63, and 0.78. The presence of correlated high-variance components in both lipidomic and transcriptomic data points to an integrated molecular system. The observation that dominant spatial gradients are conserved across both layers suggests a coordinated relationship that defines the brain's primary organizational axes.

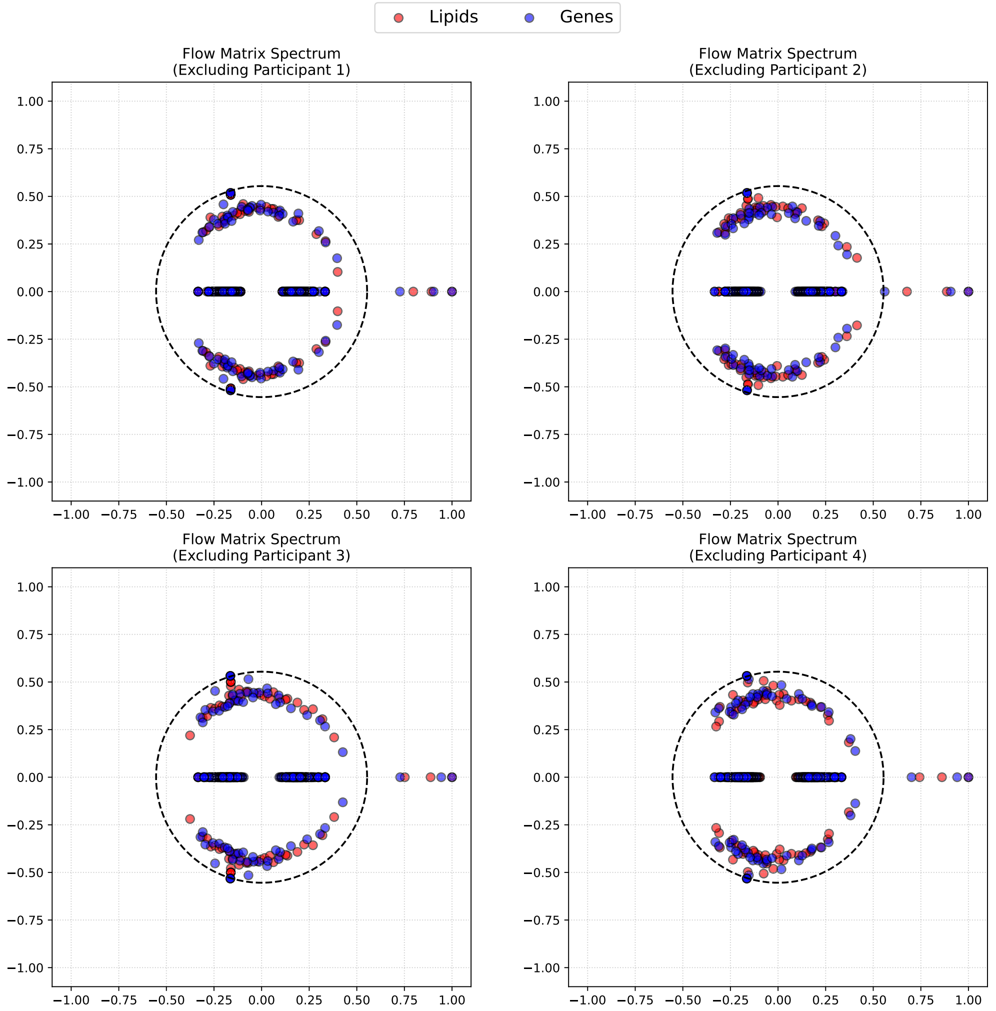


Supplementary Figure 5: Robust Community Structure in Brain Molecular Networks Revealed by Flow Matrix Spectral Analysis. This figure displays the eigenvalue spectra of the flow matrices for both transcriptomic (blue) and lipidomic (red) brain region networks, with each panel corresponding to a leave-one-out cross-validation experiment. The underlying networks were constructed using a k-Nearest Neighbors (k-NN) approach, where k was dynamically chosen for each dataset as the minimum value required to ensure full graph connectivity. The flow matrix, a variant of the non-backtracking matrix, is designed to reveal community structure through its spectral properties. The dashed black circle in each plot represents the theoretical boundary of the spectral bulk for a random graph of equivalent degree. Eigenvalues located outside this bulk in the complex plane are indicative of significant, non-random community structure. Across all four leave-one-out experiments, and for both the transcriptome and lipidome, the analysis consistently reveals three isolated eigenvalues in the tail of the distribution, lying distinctly outside the spectral bulk. The presence of these three outliers signifies the existence of three stable communities within the brain's molecular networks.

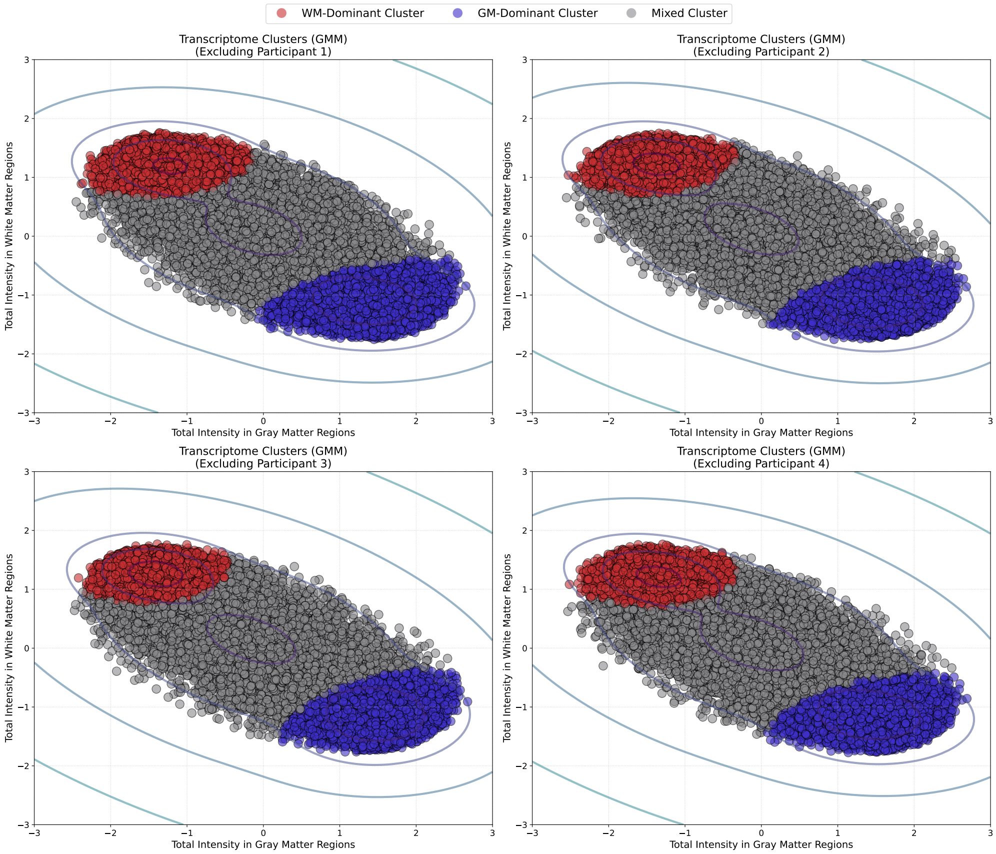


Supplementary Figure 6: Stability of Transcriptomic Clusters Assessed by Gaussian Mixture Modeling. This figure displays the results of a Gaussian Mixture Model (GMM) clustering applied to the brain's transcriptome, with each panel representing a leave-one-out cross-validation experiment where one of four different participants was excluded. Each of the 15013 genes is plotted as a point in a 2D space defined by its total summed expression in gray matter regions (x-axis) and white matter regions (y-axis). The GMM analysis identifies three distinct functional classes of genes, which are colored consistently across all four experiments: a white matter-dominant cluster (red), a gray matter-dominant cluster (blue), and a mixed-profile cluster (gray). The overlaid contour lines represent the negative log-likelihood of the GMM. The stability of the three transcriptomic clusters in terms of size, position, and density was confirmed across all leave-one-out experiments.

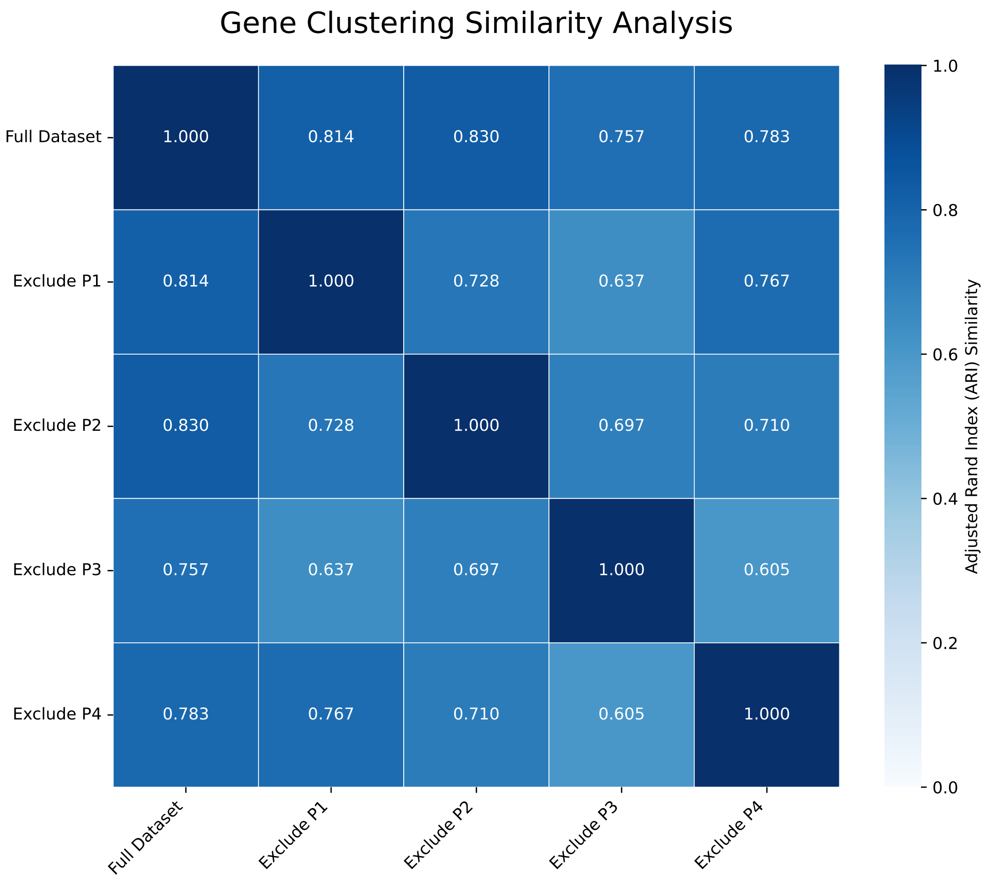


Supplementary Figure 7: Quantitative Assessment of Transcriptomic Cluster Stability via Leave-One-Out Cross-Validation. This figure presents a 5x5 heatmap quantifying the similarity between the transcriptomic cluster assignments across different datasets. The analysis compares the three functional gene clusters (WM-dominant, GM-dominant, Mixed) derived from the full dataset against those derived from four independent "leave-one-out" experiments, where a different participant was excluded from each. Similarity is measured using the Adjusted Rand Index (ARI), a metric that assesses the agreement between two clusterings while correcting for chance. An ARI of 1.0 indicates perfect agreement, while an ARI near 0 suggests a random assignment. The annotated values in each cell represent the precise ARI score for that pairwise comparison. The consistently high ARI scores demonstrate the robustness of the transcriptomic clustering. The clustering solution from the full dataset shows agreement with each of the leave-one-out experiments (ARI values of 0.814, 0.830, 0.757, and 0.783).

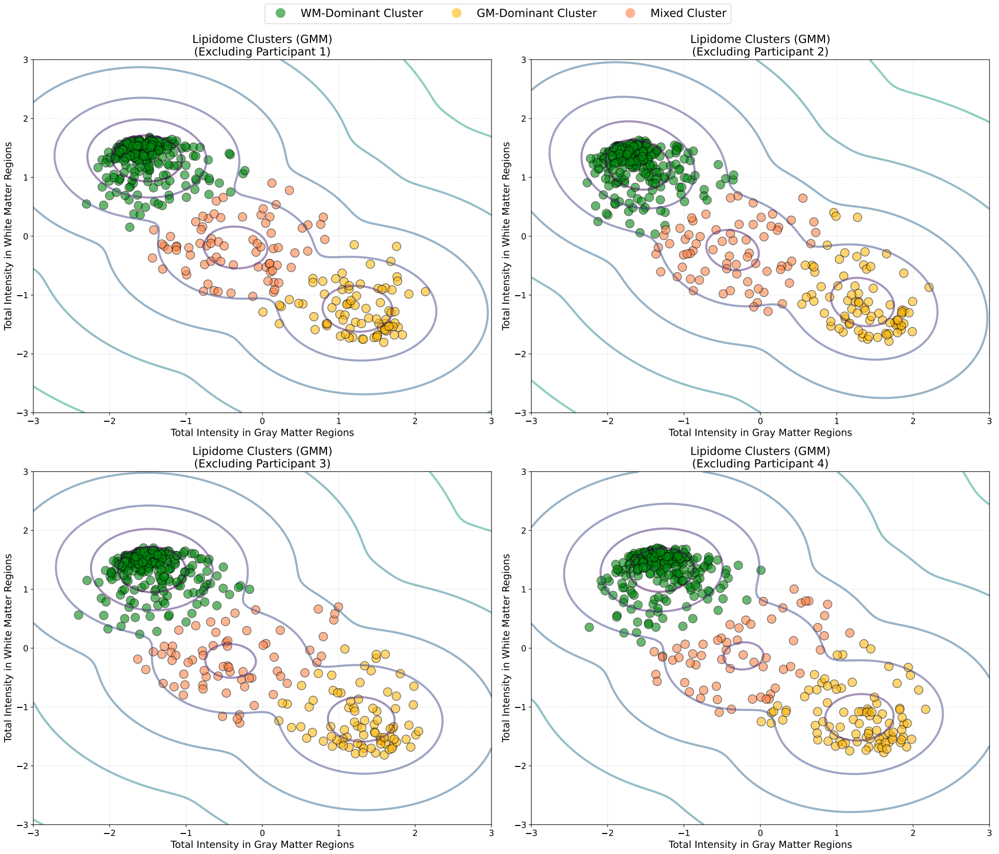


Supplementary Figure 8: Stability of Lipidomic Clusters Assessed by Gaussian Mixture Modeling. This figure displays the results of a Gaussian Mixture Model (GMM) clustering applied to the brain's lipidome, with each panel representing a leave-one-out cross-validation experiment where one of four different participants was excluded. Each of the 419 lipids is plotted as a point in a 2D space defined by its total summed concentration in gray matter regions (x-axis) and white matter regions (y-axis). The GMM analysis identifies three distinct functional classes of lipids, which are colored consistently across all four experiments: a white matter-dominant cluster (green), a gray matter-dominant cluster (yellow), and a mixed-profile cluster (orange). The overlaid contour lines represent the negative log-likelihood of the GMM, illustrating the model-derived boundaries between these functional groupings. The consistent structure of the three lipid clusters across all four leave-one-out experiments indicates a stable underlying lipidomic organization.

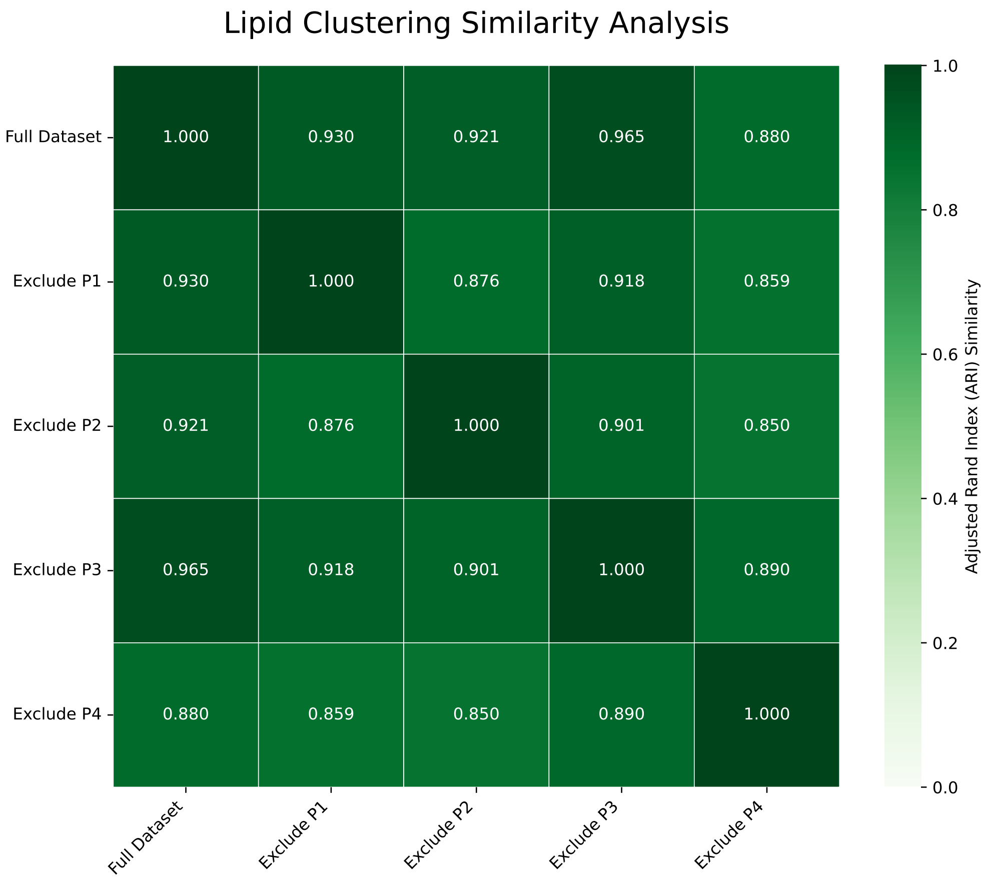


Supplementary Figure 9: Quantitative Assessment of Lipidomic Cluster Stability via Leave-One-Out Cross-Validation. This figure presents a 5x5 heatmap quantifying the similarity between the lipidomic cluster assignments across different datasets. The analysis compares the three functional lipid clusters (WM-dominant, GM-dominant, Mixed) derived from the full dataset against those derived from four independent "leave-one-out" experiments, where a different participant was excluded from each. Similarity is measured using the Adjusted Rand Index (ARI), a metric that assesses the agreement between two clusterings while correcting for chance. An ARI of 1.0 indicates perfect agreement, while an ARI near 0 suggests a random assignment. The annotated values in each cell represent the precise ARI score for that pairwise comparison. The consistently high ARI scores demonstrate the robustness of the lipidomic clustering, mirroring the stability observed in the transcriptome. The clustering solution from the full dataset shows agreement with each of the leave-one-out experiments (ARI values of 0.930, 0.921, 0.965, and 0.880), indicating that the removal of any single participant's data does not substantially alter the overall classification of lipids.

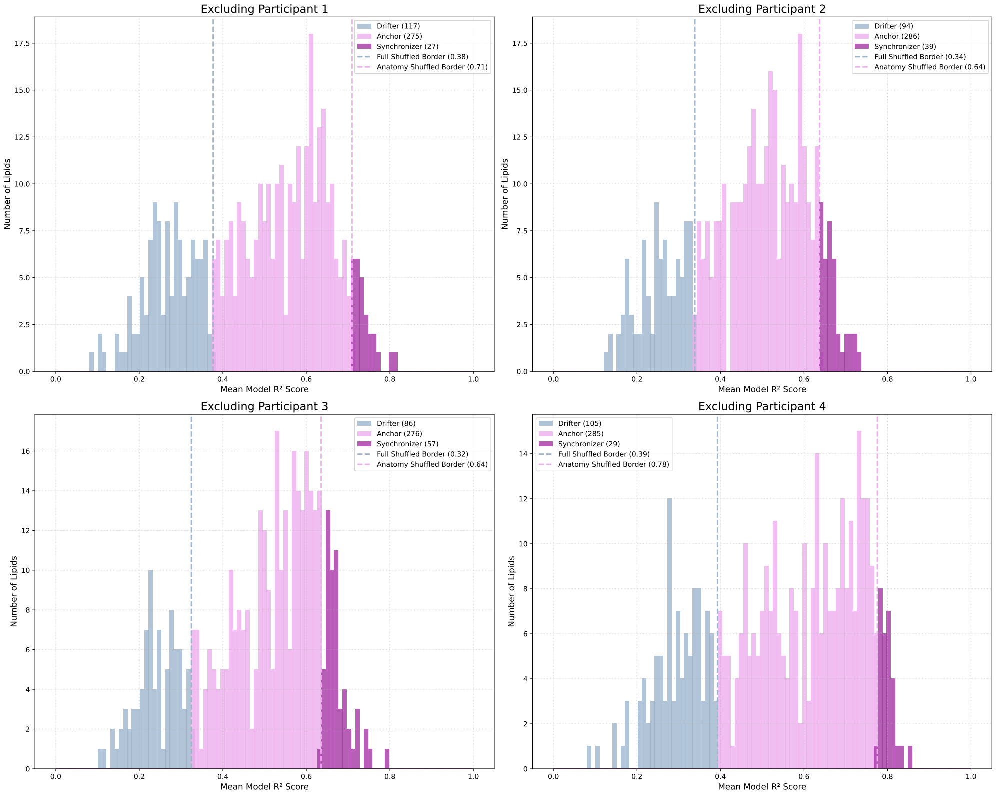


Supplementary Figure 10: Functional Categorization of Lipids Based on Predictive Modeling Performance Across Leave-One-Out Experiments. This figure presents a 2x2 grid of histograms, with each panel depicting the distribution of predictive model performance (mean $r^2$ scores) for all 419 lipids in a different leave-one-out cross-validation experiments. Each lipid is assigned to one of three functional categories based on its $r^2$ value relative to statistical null models: **Drifters** (light blue), with $r^2$ below the significance threshold of a fully shuffled model; **Anchors** (light pink), with $r^2$ between the fully shuffled and an anatomy-preserving shuffled model; and **Synchronizers** (purple), with $r^2$ exceeding both null models. The consistent partitioning of the lipidome into these three distinct classes across all four independent experiments demonstrates the stability and robustness of this functional classification framework.

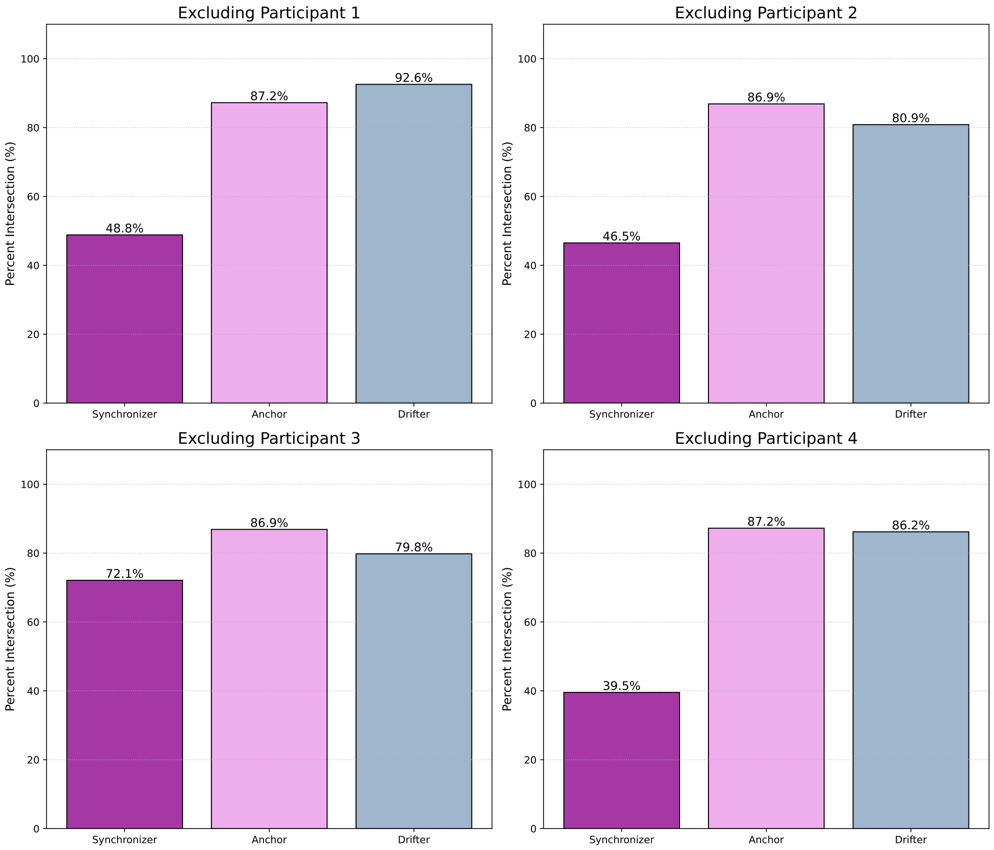


Supplementary Figure 11: Quantitative Stability of Lipid Functional Categories Across Leave-One-Out Experiments. This figure presents a 2x2 grid of bar plots, with each panel quantifying the stability of the three functional lipid categories (Synchronizers, Anchors, Drifters) for a different leave-one-out experiment. Stability is measured as the percent intersection, which is the percentage of lipids assigned to a specific category in the full dataset (ground truth) that retain the same category assignment in the leave-one-out analysis. Each bar represents one of the three categories, color-coded for consistency: Synchronizers (purple), Anchors (light pink), and Drifters (light blue). The analysis reveals stability for the Anchor and Drifter categories. The Anchor category, in particular, shows an intersection of approximately 87% across all four experiments (87.2%, 86.9%, 86.9%, and 87.2%). The Drifter category is also stable, with intersection percentages of 92.6%, 80.9%, 79.8%, and 86.2%. In contrast, the Synchronizer category (the smallest), while still showing substantial overlap, exhibits greater variability across the experiments (48.8%, 46.5%, 72.1%, and 39.5%).

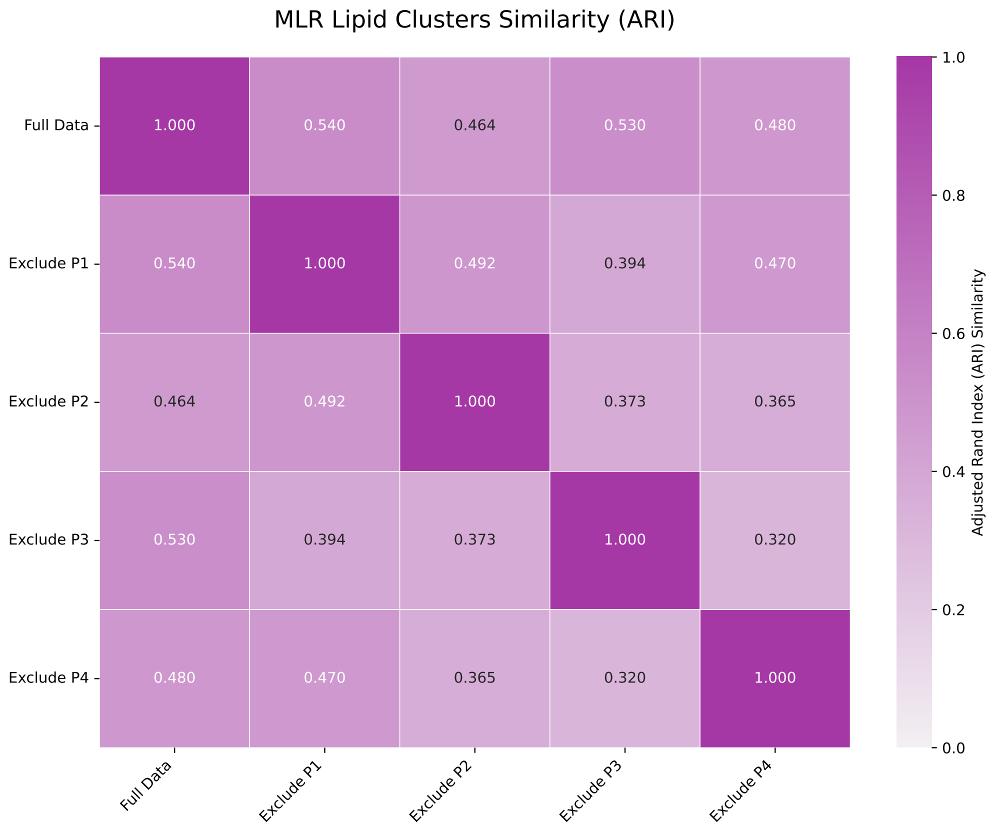


Supplementary Figure 12: Stability Assessment of the MLR-Based Lipid Functional Categories. This figure displays a 5x5 heatmap that quantifies the similarity between the MLR-derived lipid functional categories (Drifters, Anchors, Synchronizers) across the full dataset and the four leave-one-out experiments. Similarity is calculated using the Adjusted Rand Index (ARI), a metric that measures the agreement between two sets of categorical labels while correcting for chance. The annotated values represent the precise ARI score for each pairwise comparison, with 1.0 indicating identical classifications. The analysis reveals moderate to high stability in the lipid categorization. The classification from the full dataset shows agreement with those from the leave-one-out experiments, yielding ARI values of 0.540, 0.464, 0.530, and 0.480. This demonstrates that while a core organizational structure is preserved, the classification of a subset of lipids is sensitive to the removal of individual participant data.

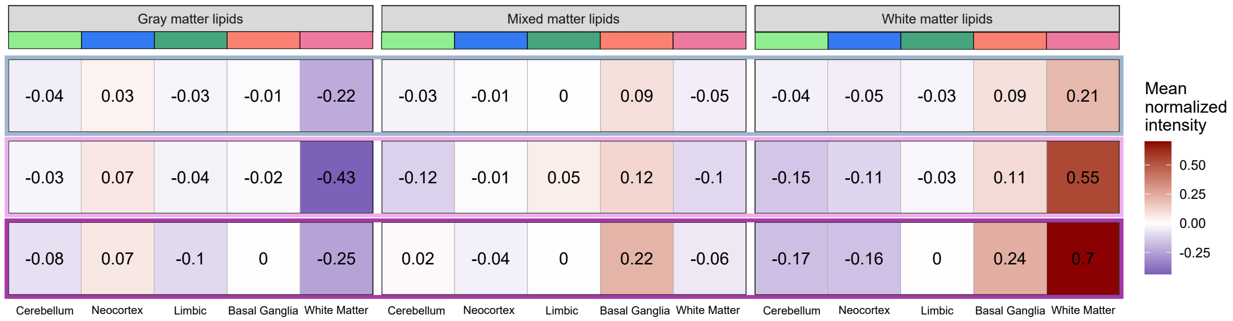


Supplementary Figure 13: Anatomical Distribution of Lipid Functional Classes. This figure details the mean normalized intensity of lipid groups across five major brain systems. The visualized lipid groups are defined by the three-way intersection of functional classes derived from MLR analysis (Synchronizers, Anchors, Drifters), anatomical clusters from GMM analysis (White Matter-Enriched, Gray Matter-Enriched, and Mixed), and the five principal brain systems (cerebellar gray matter, neocortex, limbic system, basal ganglia, and white matter). Each bar represents the average expression for a specific subgroup, revealing the precise anatomical enrichment patterns for each of the nine distinct lipid classifications, such as "White-Matter Synchronizers" or "Gray-Matter Drifters," within each colored brain system. The quantitative data reveal distinct signatures for each functional class. Synchronizers show the strongest and most consistent pattern, with White-Matter Synchronizers being massively enriched in white matter (intensity of 0.70). This highlights their coupling to gene expression programs specific to myelinated tracts. Notably, Synchronizers also show a moderate enrichment in the basal ganglia (intensities of 0.22 and 0.24 for Mixed and White-Matter classes), suggesting a secondary, but significant, transcriptional coupling to processes specific to this subcortex brain regions. Anchors perfectly illustrate their anatomy-constrained nature: Gray-Matter Anchors are strongly depleted in white matter (-0.43), while White-Matter Anchors are strongly enriched (0.55). This demonstrates that their distribution is faithfully anchored to the broad white-gray matter division. Finally, Drifters lack a clear, overarching anatomical signature; their intensity values are moderate and mixed, with the most extreme values being a mild enrichment in white matter (0.21) and a mild depletion (-0.22). This subdued and inconsistent profile supports their classification as lipids that are largely decoupled from both transcriptional programs and major anatomical structures.
